# Neonatal brain volumes and birth characteristics predict behavioural outcomes in toddlerhood

**DOI:** 10.1101/2025.11.01.686012

**Authors:** Yumnah T. Khan, Alex Tsompanidis, Carrie Allison, Richard A.I. Bethlehem, Simon Baron-Cohen

**Affiliations:** Autism Research Centre, Department of Psychiatry, University of Cambridge, Cambridge, UK, CB2 8AH; Department of Psychology, University of Cambridge, Cambridge, CB2 3EB, UK

**Keywords:** brain structure, birth factors, early behaviour, sex differences, autism

## Abstract

**Background:** Early brain structure and birth factors (e.g., sex, birth weight, gestational age at birth) are understood as critical to shaping lifelong developmental and psychopathological outcomes.

**Methods:** Using data from the Developing Human Connectome Project, we examined whether neonatal brain volumes and birth factors predict developmental and socioemotional outcomes in toddlerhood. Structural MRI scans were acquired from 391 infants at birth (193 females, 198 males; mean age = 8 days), with follow-up behavioural assessments conducted in toddlerhood (mean age = 18 months).

**Results:** Results demonstrated that larger neonatal brain volumes were associated with lower autistic traits and higher cognitive, language and motor outcomes. A higher gestational age and weight at birth were associated with higher scores on various of these outcomes - an effect that was partially mediated by larger brain volumes at birth. Females showed higher language scores compared to males, though this effect was suppressed rather than mediated by neonatal brain volumes.

**Conclusions:** These findings demonstrate how birth factors interactively shape early developmental and neuropsychiatric outcomes.

## INTRODUCTION

The prenatal and early postnatal periods are critical phases in human brain development. During these periods, the brain undergoes structural growth with remarkable speed and complexity, laying the foundation for subsequent brain development and function. Evidence is accumulating for the significance of this early period in predicting lifelong cognitive, behavioural, developmental, and health outcomes (1–4). However, while brain-behaviour associations have been predominantly researched during later stages of development, these links remain comparatively less characterised during the earliest stages of life – a period during which both brain structure and behaviour undergo rapid and dynamic changes (5–7).

Existing research, predominantly in infants born preterm, has shown that larger brain volumes at birth are associated with higher scores on a range of childhood behavioural and cognitive outcomes. For instance, larger neonatal cerebellar volumes have been linked to higher scores on psychomotor functioning (8) and autistic traits (9); larger hippocampal volumes with reduced hyperactivity, emotional, behavioural and peer problems (10); and larger thalamic and basal ganglion volumes with higher scores on motor, intelligence, and academic measures (11). However, the majority of existing studies have focused predominantly on preterm or clinical populations. Studying broader, population-level variability is critical to building a more comprehensive and generalisable picture of how early life biology shapes the course of development.

Beyond brain structure, broader birth factors, such as gestational age and weight at birth, are also robust predictors of future developmental outcomes. While the effects of preterm birth are well-documented, studies suggest that even within the term range, variations in gestational age can impact future outcomes (12). For instance, being born at a later gestational age has been linked to higher scores on motor, cognitive, language, academic, and mental health measures (13,14). Similarly, birth weight has been positively associated with cognitive, neurodevelopmental, and psychopathological outcomes, even within the typical birth weight range (Cortese et al., 2021; Gonçalves et al., 2024.; Pettersson et al., 2019). It is important to note, however, that both gestational age at birth and birth weight are strongly associated with neonatal brain volumes (18). As such, it is important to distinguish whether these factors are directly linked to developmental outcomes, or whether their effects are mediated indirectly via brain structure (i.e., a smaller gestational age/weight at birth impacts brain structure, which, in turn, impacts future outcomes).

Sex at birth is another important biological factor that affects both brain development as well as a range of outcomes across the lifespan. Studies have shown that sex differences in brain volumes are present from birth (19), with males and females showing diverging growth trajectories from prenatal development (7). These neuroanatomical differences are paralleled by sex differences in behaviour and cognition, which have also been observed as early as birth (20). During infancy, differences also begin to emerge in motor, language, and cognitive outcomes, with females generally showing faster maturation compared to males (21–24). For instance, males, on average, are more likely to take longer to develop language skills (25) and aspects of social cognition, such as joint attention (26). They are also more likely to have higher autistic traits compared to females, as measured in parent-report questionnaires (27). While neonatal brain volume has recently been shown to associate negatively with autistic traits (28), the role of baseline sex differences in this effect remains unclear. Additionally, while sex differences in both the brain and behaviour are fairly well-established, the link between the two is less understood in early life. Since differences are concurrently present across both domains, it is possible that sex differences in behavioural outcomes can be partially explained by early emerging differences in brain structure.

To understand how perinatal growth predicts future cognitive development and behaviour, the present study aimed to: (a) investigate whether neonatal brain structure predicts a range of behavioural and cognitive outcomes in toddlerhood (e.g., motor, language, and cognitive development, internalising and externalising traits, and autistic traits), (b) examine whether neonatal brain volumes mediate the associations between birth factors (e.g., gestational age, birth weight) and these outcomes, and (c) explore whether sex differences in early brain structure mediate sex differences in future outcomes. To address these objectives, we leveraged data from the developing Human Connectome Project (dHCP) (29), where infants underwent structural MRI shortly after birth followed by behavioural assessments in toddlerhood.

## METHODS AND MATERIALS

### Participants

Participants were recruited as part of the developing Human Connectome Project (dHCP) (29). Structural MRI data were acquired neonatally (mean age = 8.55 days), and follow-up behavioural assessments were conducted in toddlerhood (mean age = 18.98 months). The exclusion criteria employed in this study included preterm births (< 37 weeks gestational age), multiple births, the presence of brain anomalies in the MRI scan with likely analytical and clinical significance, pregnancy or neonatal clinical complications, and a postnatal age > 28 days at the time of the MRI scan to capture the neonatal period. Participants with any missing behavioural measures at follow-up were also excluded. The final sample consisted of 391 infants (193 females, 198 males), and further sample characteristics are reported in Table 1. As assessed by two-sample t-tests, no sex differences were observed in any of the sample characteristics.

**Table 1.** Sample Demographic Characteristics. Values represent means and, in brackets, standard deviations. ^a^ *p*-values reflect sex differences assessed by independent-samples *t*-tests.

|  | Mean (SD) -<br>All | Mean (SD) -<br>Males | Mean (SD) -<br>Females | *p*<sup>a</sup> |
| --- | --- | --- | --- | --- |
| **Gestational age at scan (weeks)** | 41.39 (1.64) | 41.23 (1.61) | 41.55 (1.66) | 0.270 |
| **Postnatal age at scan (days)** | 8.55 (8.10) | 7.88 (7.72) | 9.25 (8.44) | 0.095 |
| **Age at follow-up (months)** | 18.98 (1.99) | 18.84 (2.02) | 19.11 (1.95) | 0.162 |
| **Gestational age at birth (weeks)** | 40.17 (1.15) | 40.11 (1.15) | 40.24 (1.15) | 0.270 |
| **Birth weight (kilograms)** | 3.44 (0.47) | 3.48 (0.45) | 3.40 (0.49) | 0.089 |
| **Maternal age at birth (years)** | 33.84 (4.67) | 33.91 (4.40) | 33.77 (4.95) | 0.756 |
| **Paternal age at birth (years)** | 36.18 (5.94) | 36.22 (6.10) | 36.14 (5.78) | 0.816 |

### Outcome Measures

#### Q-CHAT

The Quantitative Checklist for Autism in Toddlers (Q-CHAT) is an 18-item parent-report questionnaire designed to quantitively measure autistic traits in toddlers through assessing various social-communication, repetitive, stereotyped, and sensory behaviours (27). While not a diagnostic instrument, it aims to capture variations in autistic traits, which are understood to exist along a continuum within the general population.

#### The Bayley Scales of Infant and Toddler Development, Third Edition (Bayley-III)

*Bayley-III* is a standardised assessment, administered by a trained professional, that evaluates infant development through structured tasks and observations (30). The assessment provides age-normed scores across cognitive, language (receptive and expressive), and motor (gross and fine) domains.

#### Child Behavioural Checklist (CBCL) for Ages 1.5–5

The CBCL is a 100-item questionnaire measuring the frequency of behavioural and emotional problems in young children (31). The measure contains problem behaviour syndrome subscales which can be grouped into two higher-order factors: internalising (measuring emotionally reactive, anxious/depressed, somatic, and withdrawn traits) and externalising (measuring attention problems and aggressive behaviour).

### MRI Data Acquisition

Anatomical data acquisition parameters are specified in the dHCP protocol (29). Data were collected using a 3-Tesla Philips Achieva system (Philips Medical Systems) with the dHCP neonatal brain imaging system, which included a neonatal 32 channel phased array head coil and a customised patient handling system (Rapid Biomedical GmbH, Rimpar, Germany). Infants were scanned without sedation after being fed and swaddled. Earplugs (President Putty, Coltene Whaledent, Mahwah, NJ, USA) and neonatal earmuffs (MiniMuffs, Natus Medical Inc., San Carlos, CA, USA) were used for auditory protection. Heart rate, oxygen saturation, and temperature were monitored throughout the scans.

The imaging protocol was designed to maximise contrast-to-noise ratio by using a Cramer Rao Lower bound approach (32). Both T2-weighted and T1-weighted inversion recovery Fast Spin Echo (FSE) images were obtained in sagittal and axial planes with relaxation times set at T1/T2: 1800/150ms for gray matter and at T1/T2: 2500/250 ms for white matter (33). The in-plane resolution was 0.8 × 0.8 mm^2^, with a slice thickness of 1.6 mm and an inter-slice overlap of 0.8 mm; for T1-weighted sagittal images, the overlap was 0.74 mm. Specific sequence parameters included: for T2-weighted images, a repetition time (TR) of 12000 ms, echo time (TE) of 156 ms, and SENSE acceleration factors of 2.11 (axial) and 2.60 (sagittal); for T1-weighted images, TR/TI/TE values were 4795/1740/8.7 ms with SENSE factors of 2.27 (axial) and 2.66 (sagittal). In addition, 3D MPRAGE images were collected at an isotropic resolution of 0.8 mm, with TR/TI/TE of 11/1400/4.6 ms and a SENSE factor of 1.2 in the right-left direction. To enable high-resolution volumetric analysis, images were processed with motion correction algorithms, and axial and sagittal scans were combined into a single 3D image for high resolution and accurate segmentation (34).

### MRI Data Pre-Processing

Structural MRI data were pre-processed using the dHCP pipeline (35). T2-weighted images underwent motion correction, bias field correction, and brain extraction using the Brain Extraction Tool (Smith, 2002). A probabilistic tissue atlas was then aligned to the bias-corrected T2-weighted scans, and initial classification of tissue types (cerebrospinal fluid, white matter, cortical grey matter, and subcortical grey matter) was carried out using the Draw-EM segmentation algorithm (37). Labelled brain atlases (38) were registered to each subject’s images using multi-channel registration that incorporated both T2-weighted image intensities and gray matter probability maps derived from the initial segmentation. This process generated a detailed segmentation encompassing 87 grey and white matter regions (37–39).

### Statistical Analysis

For each behavioural measure, associations between gray matter volumes at birth and outcomes in toddlerhood were assessed using linear regression models. Covariates included sex, postconceptional age at the neonatal scan, and infant age at follow-up (with the exception of Bayley-III scores, which were already standardised based on the toddler’s age). These models were applied across all brain volumes and outcomes of interest. To isolate regional effects independent of overall brain size, additional models were ran using total brain volume as a covariate.

Sex differences in behavioural outcomes were assessed using two-sample t-tests (for Bayley-III scores) or ANOVAs (for all remaining measures) while controlling for age at follow-up. Associations with birth weight and gestational age at birth were examined using linear regressions. Mediation analyses (bootstrapping with 5,000 resamples and bias-corrected 95% confidence intervals) were then conducted to test whether neonatal brain volumes accounted for any observed associations. In these models, sex, gestational age, or birth weight served as predictors, behavioural measures as outcomes, and pre-selected neonatal brain volumes (which showed significant associations with both the predictor and outcome) as mediators (see Figure 1). Gestational age at scan was included as a covariate in all models, while total brain volume was included as a covariate in models where regional effects, independent of total brain size, were identified. False discovery rate (FDR) corrections (40) were applied at a threshold of 0.05 within the following sets of analyses: (a) models assessing brain-behaviour associations, separately for each outcome, (b) all models assessing associations between gestational age at birth and outcomes, (c) all models assessing associations between birth weight and outcomes, (d) all models assessing sex differences in outcomes, and (e) mediation models, separately for each outcome.

**Figure 1.**
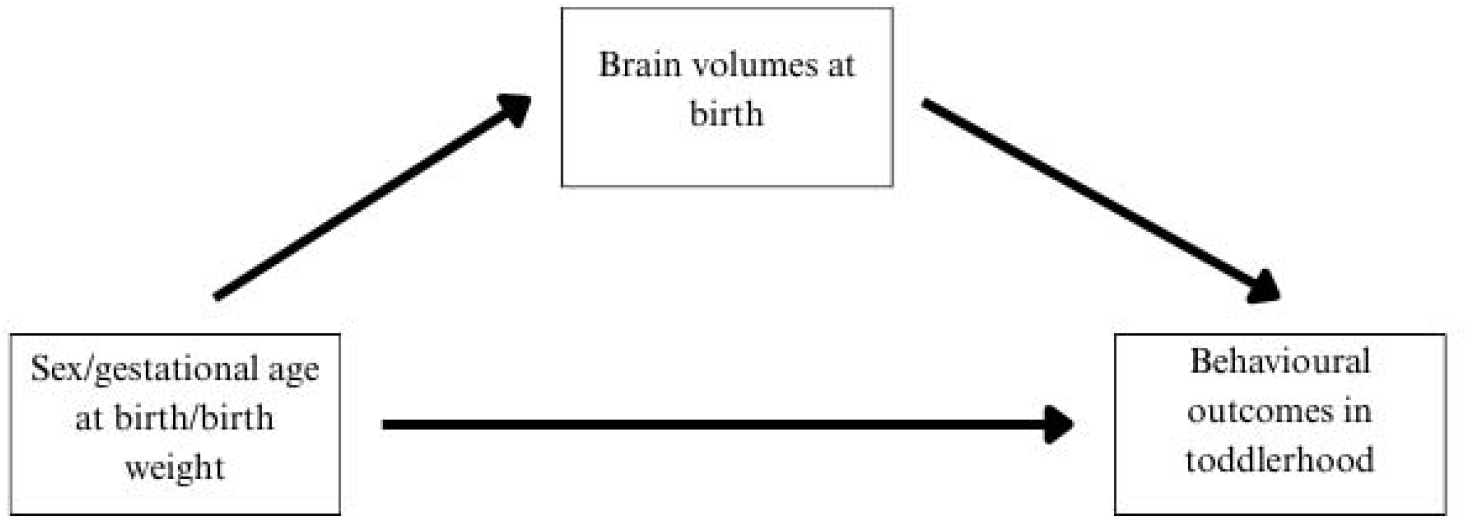
Mediation model.

## RESULTS

### Correlations between behavioural outcomes

Table 2 presents pairwise correlations between all outcome measures at 18 months. Motor, cognitive, and language scores all showed moderate associations with one another. Higher Q-CHAT scores were associated with higher externalising and internalising traits, and with lower cognitive, language, and motor scores. Internalising showed negative associations with cognitive, language, and motor outcomes, while externalising showed no significant associations with these outcomes.

**Table 2.** Pairwise correlations between behavioural outcomes at age 18 months. Correlation matrix displaying associations between behavioural outcomes at 18 months. Values represent Pearson correlation coefficients and asterisks indicate significance levels (\**p* <.05, \*\**p* <.01, \*\*\**p* <.001).

|  | Q-CHAT | Externalising | Internalising | Cognitive | Language | Motor |
| --- | --- | --- | --- | --- | --- | --- |
| **Q-CHAT** | - | **0.21** *** | **0.39** *** | **-0.32** *** | **-0.49** *** | **-0.22** *** |
| **Externalising** | **0.21** *** | - | **0.54** *** | -0.00 | -0.06 | -0.00 |
| **Internalising** | **0.39** *** | **0.54** *** | - | **-0.21** *** | **-0.20** *** | **-0.12** * |
| **Cognitive** | **-0.32** *** | -0.00 | **-0.21** *** | - | **0.59** *** | **0.53** *** |
| **Language** | **-0.49** *** | -0.06 | **-0.20** *** | **0.59** *** | - | **0.48** *** |
| **Motor** | **-0.22** *** | -0.00 | **-0.12** * | **0.53** *** | **0.48** *** | - |

#### Autistic traits

Higher scores on the Q-CHAT were associated with lower total brain, total subcortical grey matter, and total white matter volumes. Regionally, higher scores were associated with lower grey matter volumes in the bilateral thalamus, right lentiform nucleus, bilateral posterior parahippocampal gyri, brainstem and left anterior and bilateral posterior medial and inferior temporal gyri (Table 3, Supplementary Table 1). No significant regional associations were identified after controlling for total brain volume (Supplementary Table 2).

**Table 3.** Regional associations with autistic traits. Associations between brain volumes at birth and autistic traits at 18-month follow-up. Linear regression coefficients (β; both standardised and und), standard errors (SE), and false discovery rate–corrected *p*-values (*p*_FDR_) are reported for each region.

| Region | Unstandardised<br>$\beta$ Coefficient | Standardised<br>$\beta$ Coefficient | SE | *p*<sub>FDR</sub> |
| --- | --- | --- | --- | --- |
| Total brain volume | -0.000 | 0.000 | 0.000 | 0.027 |
| Total subcortical gray matter | -0.001 | 0.000 | 0.000 | 0.031 |
| Total white matter | -0.000 | 0.000 | 0.003 | 0.032 |
| Brainstem | -0.002 | -0.001 | 0.001 | 0.042 |
| Right thalamus | -0.003 | 0.002 | 0.001 | 0.049 |
| Left thalamus | -0.004 | 0.002 | 0.001 | 0.031 |
| Right lentiform nucleus | -0.004 | 0.001 | 0.001 | 0.031 |
| Right parahippocampal gyrus (anterior part) | -0.009 | 0.000 | 0.003 | 0.032 |
| Left parahippocampal gyrus (anterior part) | -0.008 | 0.002 | 0.003 | 0.049 |
| Medial and inferior temporal gyri (anterior part) | -0.003 | 0.001 | 0.001 | 0.036 |
| Right medial and inferior temporal gyri (posterior part) | -0.003 | 0.001 | 0.001 | 0.024 |
| Left medial and inferior temporal gyri (posterior part) | -0.003 | 0.001 | 0.001 | 0.013 |

#### Bayley-III

Higher scores on the cognitive subscale were associated with increased cortical grey matter volumes (Table 4). Regionally, higher scores were associated with increased grey matter volumes in the left thalamus, right occipital lobe, left posterior lateral occipitotemporal gyrus (fusiformis gyrus), and right medial and inferior temporal gyrus (Table 4, Supplementary Table 3). No significant regional associations emerged after controlling for total brain volume (Supplementary Table 4).

**Table 4.**
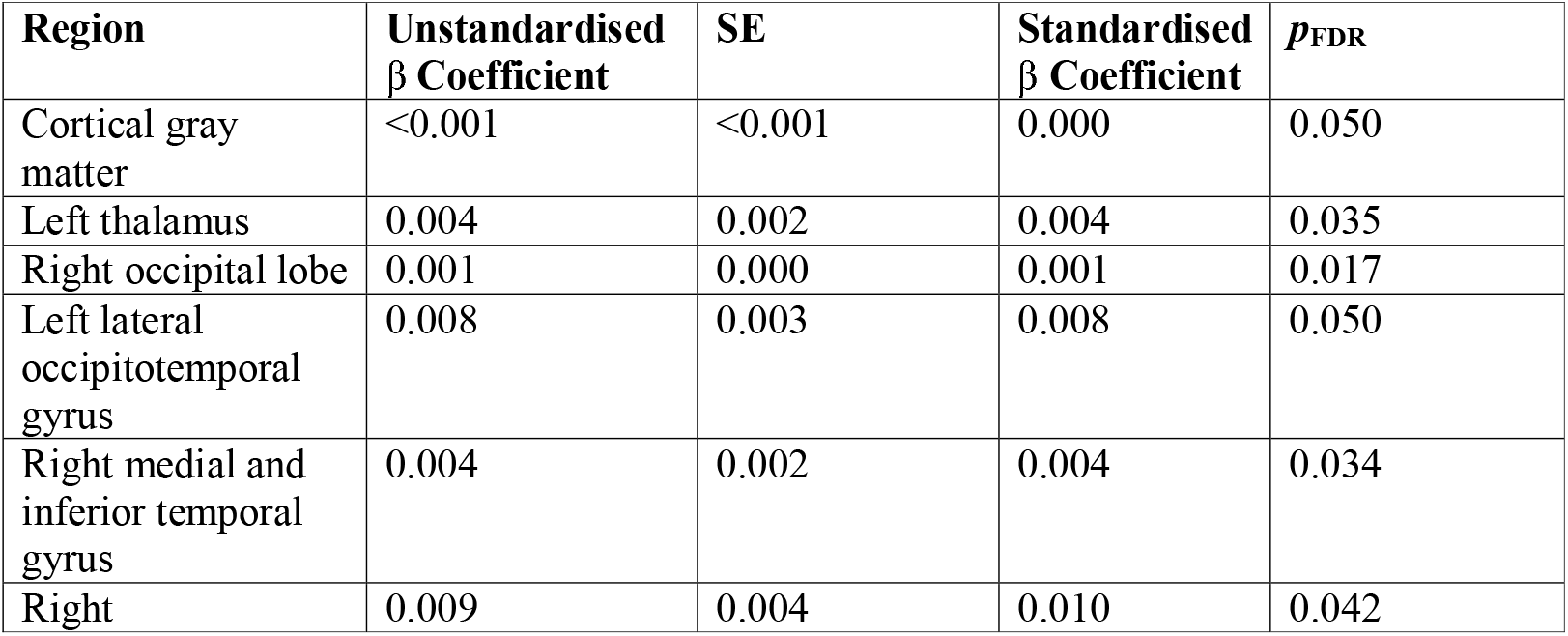
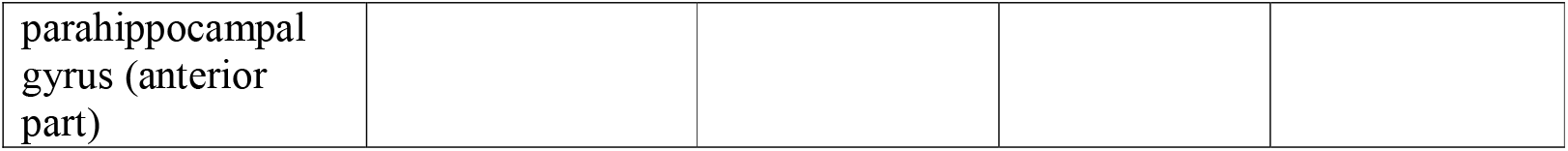
Associations with Bayley-III cognitive subscale. Associations between brain volumes at birth and cognitive outcomes at 18-month follow-up. Linear regression coefficients (β; both standardised and unstandardised), standard errors (SE), and false discovery rate–corrected *p*-values (*p*_FDR_) are reported for each region.

Higher language scores were associated with increased total brain, total white matter, and total cortical and subcortical grey matter volumes (Table 5). Regionally, higher scores were associated with increased grey matter volumes in various cortical and subcortical structures (Table 5, Supplementary Table 5). No significant regional associations were identified after controlling for total brain volume (Supplementary Table 6).

**Table 5.**
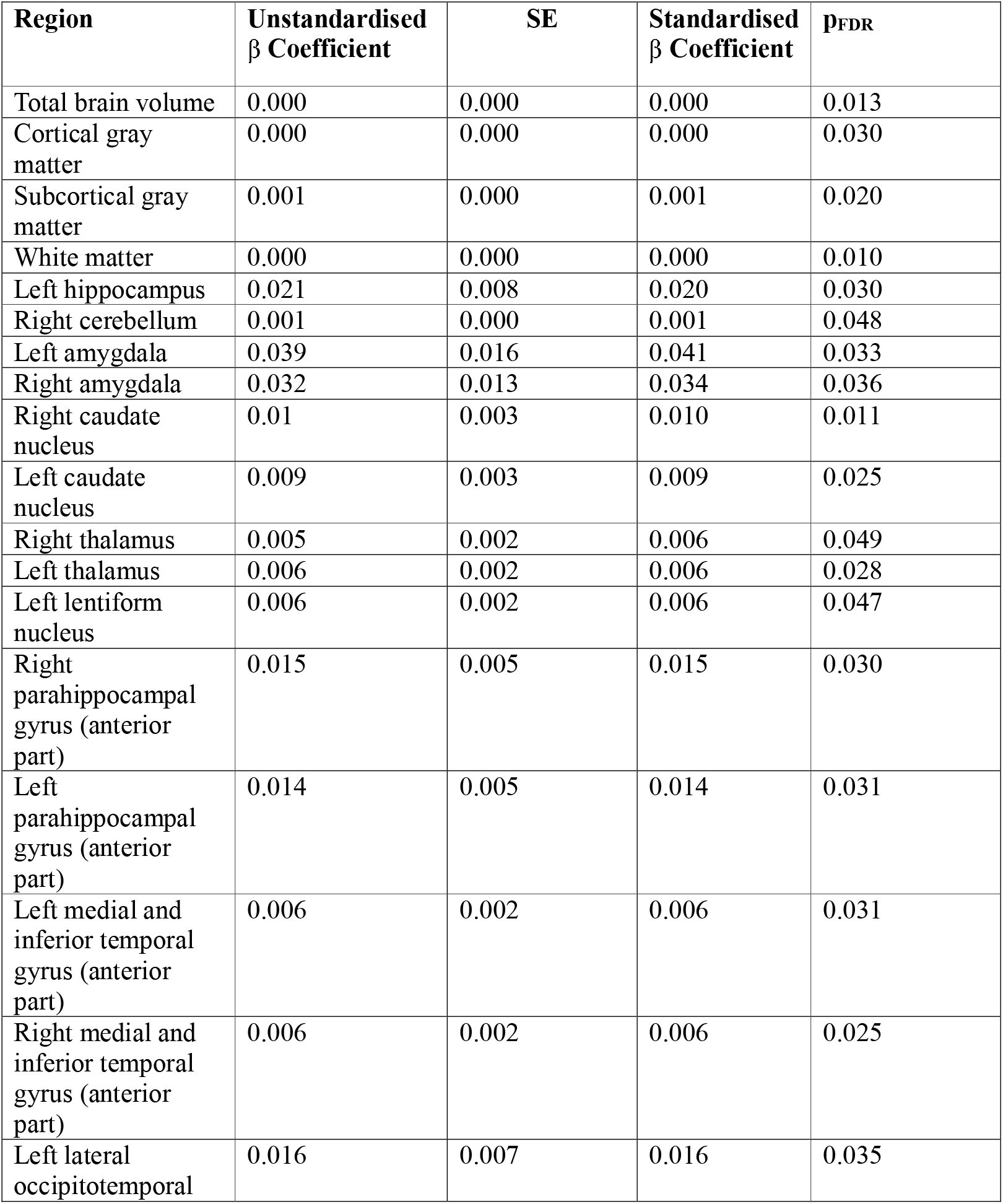
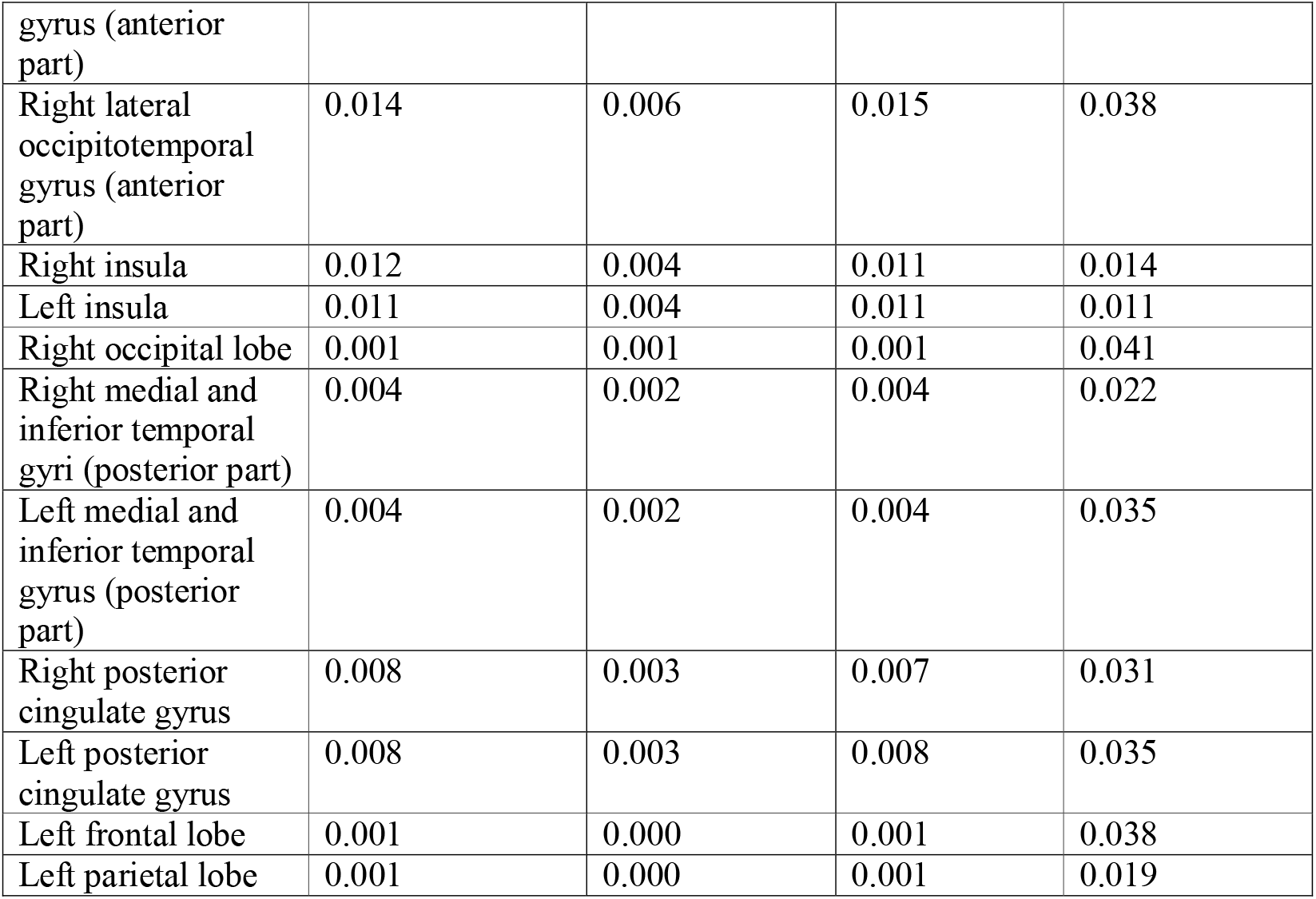
Associations with Bayley-III language subscale. Associations between brain volumes at birth and language outcomes at 18-month follow-up. Linear regression coefficients (β; both standardised and unstandardised), standard errors (SE), and false discovery rate–corrected *p*-values (*p*_FDR_) are reported for each region.

Higher motor scores were associated with decreased ventricular volumes (B= -0.001, SE = 0.000, *p*_FDR_ = 0.023) (Supplementary Table 7). After controlling for total brain volume, higher motor scores were associated with decreased volumes in the brainstem, ventricles, and left cerebellum (Table 6, Supplementary Table 8).

**Table 6.**
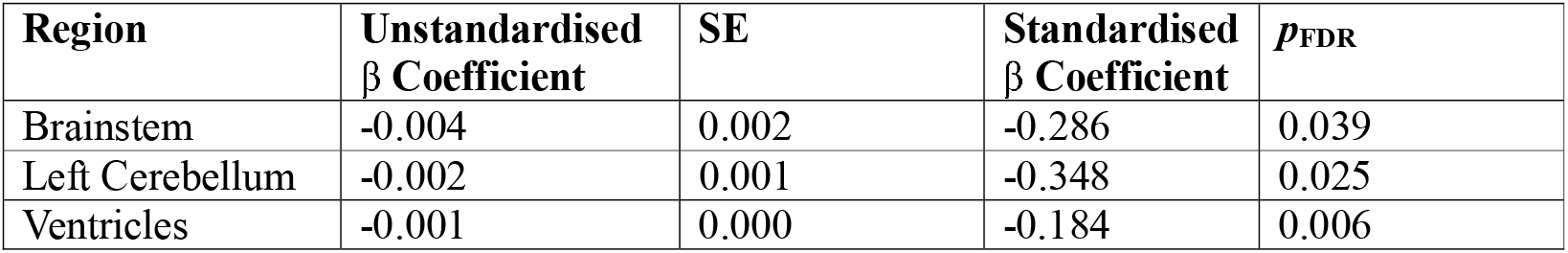
Regional associations with Bayley-III motor subscale after controlling for total brain volume. Associations between brain volumes at birth and language outcomes at 18-month follow-up after controlling for total brain volume. Linear regression coefficients (β; both standardised and unstandardised), standard errors (SE), and false discovery rate–corrected *p*-values (*p*_FDR_) are reported for each region.

| **Region** | **Unstandardised $\beta$ Coefficient** | **SE** | **Standardised $\beta$ Coefficient** | **$p_{\text{FDR}}$** |
| --- | --- | --- | --- | --- |
| Brainstem | -0.004 | 0.002 | -0.286 | 0.039 |
| Left Cerebellum | -0.002 | 0.001 | -0.348 | 0.025 |
| Ventricles | -0.001 | 0.000 | -0.184 | 0.006 |

#### CBCL

Internalising traits were negatively associated with absolute cerebellar volumes, and, after controlling for total brain volume, positively associated with frontal lobe volumes – though these associations did not survive multiple comparison corrections (Supplementary Tables 9 & 10). There were no significant associations between externalising traits and any global or regional brain volumes both before and after controlling for total brain volume (Supplementary Tables 11 & 12).

### Mediations

#### Sex Differences

Sex differences were observed in language outcomes (t = 3.44, *p*_FDR_ = 0.004, Cohen’s *d* = 0.35), with higher scores in females (M = 101.84, *SD* = 14.74) compared to males (M = 96.54, *SD* = 15.70). All regions that showed significant brain-language associations significantly *suppressed*, rather than mediated, the relationship between sex and language outcomes (Supplementary Table 13). Males showed significantly higher externalising (*F* = 3.99, *p* = 0.046, η^2^p = 0.01) and autistic traits (*F* = 4.36, *p* = 0.037, η^2^p = 0.01), though these associations did not survive multiple comparison corrections (autistic traits: *p*_FDR_ = 0.093, externalising: *p*_FDR_ = 0.093). No sex differences were observed in internalising (*F* = 0.00, *p*_FDR_ = 0.975, η^2^p = <0.001), motor (t= -0.14, *p*_FDR_ = 0.975, Cohen’s *d* = -0.01), or cognitive (t = 1.62, *p*_FDR_ = 0.161, Cohen’s *d* = 0.16) outcomes.

#### Gestational age at birth

Gestational age at birth showed significant positive associations with cognitive (β = 1.305, *p*_FDR_ = 0.021, R^2^ = 0.019), motor (β = 1.064, *p*_FDR_ = 0.021, R^2^ = 0.018), and language outcomes (β = 1.821, *p*_FDR_ = 0.021, R^2^ = 0.018). The associations with cognition were mediated by volumes in the right occipital lobe (prop. mediated = 0.575, 95% CI [0.155, 1.170], p_FDR_ = 0.024) and right parahippocampal gyrus (anterior part) (prop. mediated = 0.245, 95% CI [0.040, 0.550], p_FDR_ = 0.016). There were no significant mediations with language and motor outcomes (Supplementary Tables 14 & 15). No significant associations were identified with internalising (β = -0.193, *p*_FDR_ = 0.446, R^2^ = 0.002), externalising (β = 0.026, *p*_FDR_ = 0.933, R^2^ = 0.000), or autistic traits (β = -0.402, *p*_FDR_ = 0.422, R^2^ = 0.003).

#### Birth weight

Birth weight showed significant associations with cognitive (β = 2.527, *p*_FDR_ = 0.029, R^2^ = 0.012) and language outcomes (β = 4.019, *p*_FDR_ = 0.015, R^2^ = 0.015), while no significant associations were identified with motor (β = 1.697, *p*_FDR_ = 0.095, R^2^ = 0.007), internalising (β = -0.736, *p*_FDR_ = 0.163, R^2^ = 0.005), externalising (β = 0.784, *p*_FDR_ = 0.294, R^2^ = 0.003), or autistic traits (β = -1.571, *p*_FDR_ = 0.085, R^2^ = 0.007). The relationship between birth weight and cognition was significantly mediated by volumes in various of the structures that showed significant brain-cognition associations (Supplementary Table 16). Similarly, the relationship between birth weight and language was significantly mediated by volumes in various of the structures that showed significant brain-language associations (Supplementary Table 17).

## DISCUSSION

In the present study, we investigated the role that neonatal brain structure, sex, and perinatal factors play in predicting early cognitive and behavioural outcomes. We found that while neonatal brain structure is a reliable predictor of autistic traits, language, cognitive, and motor skills, it offers limited predictive value for psychopathological traits (i.e., internalising and externalising traits) in toddlerhood. Additionally, in some cases, neonatal brain volumes partially explain the relationship between toddlerhood outcomes and factors such as sex, weight, and gestational age at birth.

### Autistic traits

Although associations between neonatal brain structure and autistic traits have been previously been reported in the dHCP dataset (28), the present study reanalysed these data in order to (a) account for key sample differences to assess generalisability to a healthy, term-born sample (i.e., the prior study included pre-term births, which were excluded from the present research), and (b) investigate the potential role that sex and birth factors play in shaping this relationship. Consistent with the prior study, we observed that reduced brain volumes were generally associated with higher autistic traits, confirming that this association generalises to term-born samples. While autism has traditionally been associated with increased brain volumes, various explanations have been proposed to account for this observation (28). For instance, mixed findings in the literature may reflect the considerable heterogeneity within the autism spectrum, where different subtypes may follow distinct developmental trajectories. Additionally, studies have shown that autistic infants initially present with smaller head circumferences at birth followed by accelerated brain growth later in infancy, indicating that “catch-up” effects may also play a role (41).

### Motor, language, and cognitive outcomes

At the global level, larger cortical grey matter volumes were associated with higher scores on cognitive measures, replicating the well-established association between grey matter and cognition observed across various stages of development (42–44). Increased gray matter may reflect greater neuronal density, synaptic complexity, and dendritic arborisation – factors which may collectively impact cognitive processing capacities (45,46). At the regional level, consistent with prior literature, positive associations were identified with the thalamus, insula, temporal, and occipital lobes (47). These regions are broadly implicated in sensory processing and integration, and their structural maturation may facilitate early cognitive development. Language outcomes showed widespread associations with total grey and white matter, as well as with various temporal, parietal, frontal, occipital, and subcortical structures. After controlling for total brain volume, no significant associations emerged for either cognitive or language outcomes, indicating that the observed findings are primarily driven by global brain characteristics than by region-specific effects.

While motor outcomes showed no significant global or regional associations with absolute brain volumes, they were associated with reduced volumes in the brainstem and left cerebellum after controlling for total brain volume. These findings are unsurprising as both the cerebellum and brainstem play well-established roles in motor control, movement, balance, and coordination (48–50). As such, relatively increased volumes in these structures at birth may support motor development during infancy. These findings also indicate that individual differences in motor development are driven by localised regional variations rather than broad, global brain differences.

### Psychopathological outcomes

Internalising traits were negatively associated with absolute cerebellar volumes, a region frequently implicated in psychopathology (51), and positively associated with frontal lobe volumes after controlling for total brain volume. These associations did not, however, survive a correction for multiple comparisons. There were no significant associations between externalising traits and any global or regional brain volumes. It therefore appears that neonatal brain structure is a stronger predictor of developmental than psychopathological outcomes in toddlerhood. It is possible that meaningful associations may become more apparent during later stages of development (i.e., adolescence), when mental health traits become more pronounced and begin to show greater variability (52). Additionally, since mental health outcomes are heavily shaped by brain-environment interactions (53,54), associations with brain structure may be minimal prior to extensive environmental influence.

### Gestational age at birth and birth weight

As expected, our findings showed that a greater gestational age at birth was linked to improved language, cognitive, and motor outcomes. Importantly, brain volumes partially mediated these associations for cognitive outcomes, suggesting that gestational age at birth influences later development, in part, through its effects on early brain structure. Prior research has shown that, at term-equivalent age, infants born earlier within the term period have reduced brain volumes compared to those born later (18). This is because the last trimester of gestation is characterised by rapid cortical development (e.g., accelerated cortical growth, synaptogenesis, proliferation, myelination, and gyrification), marking it as a critical period for establishing the structural and functional foundations of the brain (5,6). Birth occurring even a few weeks earlier may interrupt these processes, leading to long-term alterations in brain structure and associated outcomes (13).

We additionally identified that birth weight was associated with cognitive and language outcomes, and these relationships were also mediated by neonatal brain volumes. Both birth weight and gestational age at birth are shown to have long-term developmental significance, predicting brain volumes and behavioural outcomes even later in life (Gonçalves et al., 2024.; Ma et al., 2022; Silva et al., 2021). Our findings extend prior literature by demonstrating their effects are not independent but are rather partially mediated via neonatal brain volumes. It is also worth noting that while birth factors predicted early language and cognitive outcomes, they were not significantly associated with early internalising, externalising, or autistic traits.

### Sex differences

We next examined whether sex plays a role in shaping early developmental outcomes via its effects on neonatal brain structure. Males showed higher externalising and autistic traits in toddlerhood compared to females, though these associations did not survive a correction for multiple comparisons. Females showed improved language outcomes in toddlerhood compared to males – a finding that is consistent with a well-established developmental trend in which females exhibit an on-average advantage in language, social cognition, and empathising (23,57,58). Interestingly, mediation analyses revealed that brain volumes at birth *suppressed* rather than mediated this relationship. While larger neonatal brain volumes were generally associated with improved language outcomes, females – despite on average having smaller neonatal brain volumes (19) – showed higher language scores compared to males. Although seemingly counterintuitive, this pattern can be explained by compensatory mechanisms. It has been proposed that sex differences in the brain may serve not only to create behavioural differences, but also to prevent or mitigate them by balancing hormonal or physiological factors that might otherwise amplify them further (59). According to this notion, larger brain volumes in males may act protectively to reduce a language performance gap that might otherwise be even more pronounced. It is also possible that these sex differences are better explained by metrics beyond brain volumes (e.g., functional connectivity or white matter microstructure) and vary depending on the developmental stage under investigation.

### Strengths and limitations

To the best of our knowledge, this study is among the few to examine associations between neonatal brain structure and a broad spectrum of outcomes in a large, representative, term-born sample. This prospective, longitudinal design (i.e., brain imaging at birth and outcomes assessed at 18 months) enabled us to investigate how early neuroanatomy shapes future behavioural, developmental, and socioemotional outcomes. While causality cannot be assumed, the design offers some insight into the directionality and temporal sequence of these associations. The large sample size also provides adequate statistical power to detect meaningful effects across multiple domains, while the focus on a non-clinical, term-born cohort provides insights into general, population-level variability in development.

Several limitations should also be considered. First, brain structure was assessed at a single timepoint, which fails to capture the rapid and dynamic changes that occur across the first few years of life. It is possible that infants with initially reduced brain volumes may experience relative overgrowth, while those with initially increased volumes may experience undergrowth. These developmental trajectories, rather than brain structure at a single point in time, may serve as more informative predictors of future behavioural outcomes (Vanes et al., 2023). Additionally, it is important to note that brain–behaviour associations are likely stage-specific and may vary, or become stronger, when both measurements are more temporally aligned (61). Secondly, the follow-up was limited to one timepoint in toddlerhood, and many of the studied traits continue to evolve beyond 18 months. Longitudinal studies incorporating repeated brain and behaviour measurements may provide a richer developmental account and demonstrate how changes in brain morphology over time relate to emerging behaviour. Thirdly, while we focused primarily on volumetric measures, other imaging modalities (e.g., white matter microstructure, functional connectivity) may serve as more sensitive predictors of behavioural outcomes. Finally, while perinatal factors such as gestational age and birth weight were considered, a range of additional environmental and psychosocial factors (e.g., parenting quality, maternal mental health, and socioeconomic status) were not assessed and may serve as important mediators or moderators. Future research may consider incorporating a wider array of imaging techniques and environmental variables to more comprehensively map early brain-behaviour associations.

## Conclusion

Collectively, these findings demonstrate a broad pattern in which larger neonatal brain volumes are associated with higher scores on cognitive, behavioural, psychopathological, and neurodevelopmental measures in toddlerhood. Additionally, neonatal brain structure appears to hold domain-specific predictive value, showing stronger associations with certain outcomes (e.g., language, cognition, motor and autistic traits) compared to others (e.g., internalising and externalising traits). Associations with these latter traits may become more pronounced later in development, as individual variability increases and brain–environment interactions become more influential. Our findings further demonstrate that birth factors, such as gestational age weight, influence future outcomes in part through their impact on early brain structure, reinforcing the developmental significance of late gestational development and fetal growth. Finally, sex differences may influence brain–behaviour associations through a multifaceted interplay of mediatory and compensatory mechanisms which may not be fully captured by brain volumes alone. Collectively, these findings provide further evidence that neonatal brain structure and birth factors are important markers for shaping future behavioural development.

## Acknowledgement

We would like to thank Dr Kamen Tsvetanov for providing statistical advice on the mediation models. We would also like to thank Eden Hymanson, Sehanya Wickramanayake, and Manya Gupta for assisting with organising the supplementary materials.

These results were obtained using data made available from the Developing Human Connectome Project funded by the European Research Council under the European Union’s Seventh Framework Programme (FP/2007-2013) / ERC Grant Agreement no. [319456]. Data used in the preparation of this manuscript were obtained from the National Institute of Mental Health (NIMH) Data Archive (NDA). NDA is a collaborative informatics system created by the National Institutes of Health to provide a national resource to support and accelerate research in mental health. Dataset identifier(s): ***3995***. This manuscript reflects the views of the authors and may not reflect the opinions or views of the NIH or of the Submitters submitting original data to NDA.

Y.T.K is supported by the Cambridge Trust and Trinity College, Cambridge. SBC received funding from the Wellcome Trust 214322\Z\18\Z. For the purpose of Open Access, the author has applied a CC BY public copyright licence to any Author Accepted Manuscript version arising from this submission. SBC also received funding from the Innovative Medicines Initiative 2 Joint Undertaking under grant agreement No 777394 for the project AIMS-2-TRIALS. This Joint Undertaking receives support from the European Union’s Horizon 2020 research and innovation programme and EFPIA and AUTISM SPEAKS, Autistica, SFARI. SBC also received funding from Autism Action, SFARI, the Templeton World Charitable Fund and the MRC. The funders had no role in the design of the study; in the collection, analyses, or interpretation of data; in the writing of the manuscript, or in the decision to publish the results. Any views expressed are those of the author(s) and not necessarily those of the funders (including IHI-JU2). All research at the Department of Psychiatry in the University of Cambridge is supported by the NIHR Cambridge Biomedical Research Centre (NIHR203312) and the NIHR Applied Research Collaboration East of England. The views expressed are those of the author(s) and not necessarily those of the NIHR or the Department of Health and Social Care.

