## Supplementary Materials for "Neonatal brain volumes and birth characteristics predict behavioural outcomes in toddlerhood"

Table of Contents

Supplementary Table 1: Associations with autistic traits	2

Supplementary Table 2: Associations with autistic traits after controlling for total brain volume	5

Supplementary Table 3: Associations with Bayley-III cognitive subscale	9

Supplementary Table 4: Associations with Bayley-III cognitive subscale after controlling for total brain volume	12

Supplementary Table 5: Associations with Bayley-III language subscale	16

Supplementary Table 6: Associations with Bayley-III language subscale after controlling for total brain volume	19

Supplementary Table 7: Associations with Bayley-III motor subscale	23

Supplementary Table 8: Associations with Bayley-III motor subscale after controlling for total brain volume	26

Supplementary Table 9: Associations with internalising	29

Supplementary Table 10: Associations with internalising after controlling for total brain volume	32

Supplementary Table 11: Associations with externalising	35

Supplementary Table 12: Associations with externalising after controlling for total brain volume	38

Supplementary Table 13: Brain volumes mediating the association between sex and language outcomes	41

Supplementary Table 14: Brain volumes mediating the association between gestational age at birth and language outcomes	43

Supplementary Table 15: Brain volumes mediating the association between gestational age at birth and motor outcomes after controlling for total brain volume	45

Supplementary Table 16: Regional brain volumes mediating the association between birth weight and cognitive outcomes	46

Supplementary Table 17: Regional brain volumes mediating the association between birth weight and language outcomes	47

**Supplementary Table 1. Associations with autistic traits**

Associations between brain volumes at birth and autistic traits at 18-month follow-up. Linear regression coefficients (β; both standardised and unstandardised), standard errors (SE), and false discovery rate–corrected *p*-values (*p*_FDR_) are reported for each region.

| **Region** | **Unstandardised** β **Coefficient** | **SE** | **Standardised** β **Coefficient** | ***p*_FDR_** |
| --- | --- | --- | --- | --- |
| Total brain volume | 0.000 | 0.000 | 0.000 | 0.027* |
| Total white matter | 0.000 | 0.000 | 0.000 | 0.013* |
| Total cortical grey matter | 0.000 | 0.000 | 0.000 | 0.073 |
| Total subcortical grey matter | -0.001 | 0.000 | 0.000 | 0.031* |
| Brainstem | -0.002 | 0.001 | -0.001 | 0.042* |
| Ventricles | 0.000 | 0.000 | -0.001 | 0.397 |
| CSF | 0.000 | 0.000 | 0.000 | 0.064 |
| Hippocampus left | -0.009 | 0.005 | 0.002 | 0.093 |
| Hippocampus right | -0.007 | 0.005 | 0.004 | 0.226 |
| Amygdala left | -0.014 | 0.009 | 0.014 | 0.202 |
| Amygdala right | -0.017 | 0.007 | 0.004 | 0.062 |
| Anterior temporal lobe medial part left | -0.006 | 0.003 | 0.003 | 0.117 |
| Anterior temporal lobe medial part right | -0.005 | 0.003 | 0.006 | 0.226 |
| Anterior temporal lobe lateral part left | -0.005 | 0.003 | 0.002 | 0.122 |
| Anterior temporal lobe lateral part right | -0.003 | 0.003 | 0.001 | 0.395 |
| Gyri parahippocampalis et ambiens anterior part left | -0.008 | 0.003 | 0.002 | 0.049* |
| Gyri parahippocampalis et ambiens anterior part right | -0.009 | 0.003 | 0.000 | 0.032* |
| Superior temporal gyrus middle part left | -0.002 | 0.001 | -0.001 | 0.236 |
| Superior temporal gyrus middle part right | -0.001 | 0.001 | 0.000 | 0.420 |
| Medial and inferior temporal gyri anterior part left | -0.003 | 0.001 | 0.001 | 0.036 |
| Medial and inferior temporal gyri anterior part right | -0.001 | 0.001 | 0.002 | 0.395 |
| Lateral occipitotemporal gyrus fusiformis anterior part left | -0.009 | 0.004 | -0.001 | 0.052 |
| Lateral occipitotemporal gyrus - gyrus fusiformis anterior part right | -0.006 | 0.003 | 0.003 | 0.107 |
| Cerebellum left | -0.001 | 0.000 | -0.001 | 0.062 |
| Cerebellum right | -0.001 | 0.000 | 0.000 | 0.060 |
| Brainstem | -0.002 | 0.001 | -0.001 | 0.042* |
| Insula right | -0.003 | 0.002 | 0.002 | 0.313 |
| Insula left | -0.003 | 0.002 | 0.002 | 0.185 |
| Occipital lobe right | -0.001 | 0.000 | 0.000 | 0.168 |
| Occipital lobe left | 0.000 | 0.000 | 0.000 | 0.300 |
| Gyri parahippocampalis et ambiens posterior part right | -0.002 | 0.004 | 0.001 | 0.732 |
| Gyri parahippocampalis et ambiens posterior part left | 0.002 | 0.004 | 0.000 | 0.677 |
| Lateral occipitotemporal gyrus   gyrus fusiformis posterior part right | -0.006 | 0.003 | -0.001 | 0.060 |
| Lateral occipitotemporal gyrus fusiformis posterior part left | -0.004 | 0.003 | 0.003 | 0.278 |
| Medial and inferior temporal gyri posterior part right | -0.003 | 0.001 | 0.001 | 0.024* |
| Medial and inferior temporal gyri posterior part left | -0.003 | 0.001 | 0.001 | 0.013* |
| Superior temporal gyrus posterior part right | -0.002 | 0.002 | -0.001 | 0.395 |
| Superior temporal gyrus posterior part left | -0.006 | 0.003 | 0.002 | 0.066 |
| Cingulate gyrus anterior part right | -0.003 | 0.002 | -0.001 | 0.306 |
| Cingulate gyrus anterior part left | -0.002 | 0.002 | 0.000 | 0.331 |
| Cingulate gyrus posterior part right | -0.002 | 0.002 | 0.003 | 0.244 |
| Cingulate gyrus posterior part left | -0.003 | 0.002 | 0.003 | 0.202 |
| Frontal lobe right | 0.000 | 0.000 | 0.000 | 0.117 |
| Frontal lobe left | 0.000 | 0.000 | 0.000 | 0.109 |
| Parietal lobe right | 0.000 | 0.000 | 0.000 | 0.147 |
| Parietal lobe left | 0.000 | 0.000 | 0.000 | 0.097 |
| Caudate nucleus right | -0.003 | 0.002 | 0.000 | 0.192 |
| Caudate nucleus left | -0.004 | 0.002 | 0.001 | 0.081 |
| Thalamus right | -0.003 | 0.001 | 0.002 | 0.049* |
| Thalamus left | -0.004 | 0.001 | 0.002 | 0.024* |
| Subthalamic nucleus right | -0.043 | 0.019 | 0.027 | 0.067 |
| Subthalamic nucleus left | -0.011 | 0.021 | 0.013 | 0.653 |
| Lentiform Nucleus right | -0.004 | 0.001 | 0.001 | 0.031* |
| Lentiform Nucleus left | -0.003 | 0.001 | 0.000 | 0.059 |
| Corpus Callosum | -0.001 | 0.001 | -0.001 | 0.288 |

**Supplementary Table 2. Associations with autistic traits after controlling for total brain volume**

Associations between brain volumes at birth and autistic traits at 18-month follow-up after controlling for total brain volume. Linear regression coefficients (β; both standardised and unstandardised), standard errors (SE), and false discovery rate–corrected *p*-values (*p*_FDR_) are reported for each region.

| **Region** | **Unstandardised** β **Coefficient** | **SE** | **Standardised** β **Coefficient** | **p_FDR_** |
| --- | --- | --- | --- | --- |
| Total cortical grey matter | 0.000 | 0.000 | 0.775 | 0.522 |
| Total subcortical grey matter | 0.000 | 0.001 | -0.093 | 0.669 |
| Total white matter | 0.000 | 0.000 | -0.341 | 0.566 |
| Brainstem | -0.003 | 0.003 | -0.076 | 0.342 |
| CSF | 0.000 | 0.000 | -0.077 | 0.769 |
| Ventricles | -0.001 | 0.001 | 0.005 | 0.263 |
| Hippocampus left | 0.011 | 0.010 | -0.041 | 0.424 |
| Hippocampus right | 0.005 | 0.010 | 0.007 | 0.692 |
| Amygdala left | 0.018 | 0.019 | 0.012 | 0.501 |
| Amygdala right | 0.015 | 0.016 | -0.061 | 0.505 |
| Anterior temporal lobe - medial part left | -0.001 | 0.007 | -0.040 | 0.859 |
| Anterior temporal lobe - medial part right | 0.000 | 0.007 | 0.030 | 0.989 |
| Anterior temporal lobe - lateral part left | -0.001 | 0.006 | -0.034 | 0.921 |
| Anterior temporal lobe - lateral part right | -0.005 | 0.007 | 0.069 | 0.540 |
| Gyri parahippocampalis et ambiens anterior part left | 0.007 | 0.007 | -0.071 | 0.421 |
| Gyri parahippocampalis et ambiens anterior part right | 0.007 | 0.007 | -0.086 | 0.470 |
| Superior temporal gyrus -middle part left | -0.004 | 0.003 | 0.041 | 0.368 |
| Superior temporal gyrus- middle part right | -0.004 | 0.003 | 0.137 | 0.342 |
| Medial and inferior temporal gyri anterior part left | 0.002 | 0.003 | -0.094 | 0.572 |
| Medial and inferior temporal gyri anterior part right | 0.002 | 0.003 | 0.136 | 0.540 |
| Lateral occipitotemporal gyrus - gyrus fusiformis anterior part left | 0.008 | 0.008 | -0.086 | 0.477 |
| Lateral occipitotemporal gyrus - gyrus fusiformis anterior part right | 0.006 | 0.007 | -0.029 | 0.533 |
| Cerebellum left | 0.000 | 0.001 | -0.072 | 0.714 |
| Cerebellum right | 0.000 | 0.001 | -0.084 | 0.722 |
| Brainstem | -0.003 | 0.003 | -0.076 | 0.342 |
| Insula right | 0.007 | 0.006 | 0.096 | 0.395 |
| Insula left | 0.007 | 0.005 | 0.050 | 0.349 |
| Occipital lobe right | 0.000 | 0.001 | 0.096 | 0.921 |
| Occipital lobe left | -0.001 | 0.001 | 0.185 | 0.540 |
| Gyri parahippocampalis et ambiens posterior part right | -0.007 | 0.009 | 0.140 | 0.554 |
| Gyri parahippocampalis et ambiens posterior part left | -0.009 | 0.008 | 0.154 | 0.412 |
| Lateral occipitotemporal gyrus - gyrus fusiformis posterior part right | 0.002 | 0.006 | -0.093 | 0.721 |
| Lateral occipitotemporal gyrus- gyrus fusiformis posterior part left | 0.002 | 0.006 | -0.005 | 0.714 |
| Medial and inferior temporal gyri posterior part right | 0.002 | 0.003 | -0.141 | 0.572 |
| Medial and inferior temporal gyri posterior part left | 0.001 | 0.002 | -0.173 | 0.762 |
| Superior temporal gyrus - posterior part right | -0.003 | 0.005 | 0.069 | 0.645 |
| Superior temporal gyrus - posterior part left | 0.001 | 0.005 | -0.073 | 0.811 |
| Cingulate gyrus - anterior part right | -0.004 | 0.005 | 0.063 | 0.528 |
| Cingulate gyrus - anterior part left | -0.005 | 0.004 | 0.027 | 0.404 |
| Cingulate gyrus - posterior part right | 0.002 | 0.005 | 0.094 | 0.670 |
| Cingulate gyrus - posterior part left | 0.002 | 0.005 | 0.079 | 0.709 |
| Frontal lobe right | -0.001 | 0.001 | 0.404 | 0.222 |
| Frontal lobe left | -0.001 | 0.001 | 0.398 | 0.540 |
| Parietal lobe right | -0.001 | 0.001 | 0.319 | 0.395 |
| Parietal lobe left | 0.000 | 0.001 | 0.243 | 0.747 |
| Caudate nucleus right | 0.006 | 0.004 | 0.022 | 0.282 |
| Caudate nucleus left | 0.004 | 0.004 | -0.022 | 0.495 |
| Thalamus right | 0.000 | 0.004 | -0.061 | 0.992 |
| Thalamus left | 0.002 | 0.004 | -0.132 | 0.666 |
| Subthalamic nucleus right | -0.031 | 0.044 | -0.042 | 0.572 |
| Subthalamic nucleus left | -0.056 | 0.044 | 0.080 | 0.342 |
| Lentiform Nucleus right | 0.001 | 0.003 | -0.111 | 0.837 |
| Lentiform Nucleus left | 0.001 | 0.003 | -0.058 | 0.751 |
| Corpus Callosum | -0.003 | 0.002 | 0.037 | 0.424 |

**Supplementary Table 3. Associations with Bayley-III cognitive subscale**

Associations between brain volumes at birth and cognitive outcomes at 18-month follow-up. Linear regression coefficients (β; both standardised and unstandardised), standard errors (SE), and false discovery rate–corrected *p*-values (*p*_FDR_) are reported for each region.

| **Region** | **Unstandardised** β **Coefficient** | **SE** | **Standardised** β **Coefficient** | ***p*_FDR_** |
| --- | --- | --- | --- | --- |
| Total brain volume | 0.000 | 0.000 | 0.000 | 0.077 |
| Total white matter | 0.000 | 0.000 | 0.000 | 0.122 |
| Total cortical grey matter | 0.004 | 0.002 | 0.000 | 0.032* |
| Total subcortical grey matter | 0.000 | 0.000 | 0.000 | 0.154 |
| Brainstem | 0.002 | 0.001 | 0.002 | 0.193 |
| Ventricles | -0.001 | 0.000 | -0.001 | 0.197 |
| CSF | 0.000 | 0.000 | 0.000 | 0.481 |
| Hippocampus left | 0.012 | 0.006 | 0.013 | 0.094 |
| Hippocampus right | 0.010 | 0.006 | 0.010 | 0.144 |
| Amygdala left | 0.014 | 0.011 | 0.017 | 0.258 |
| Amygdala right | 0.015 | 0.009 | 0.017 | 0.163 |
| Anterior temporal lobe medial part left | 0.001 | 0.004 | 0.001 | 0.903 |
| Anterior temporal lobe medial part right | 0.004 | 0.004 | 0.005 | 0.326 |
| Anterior temporal lobe lateral part left | 0.001 | 0.004 | 0.001 | 0.781 |
| Anterior temporal lobe lateral part right | 0.004 | 0.025 | 0.002 | 0.876 |
| Gyri parahippocampalis et ambiens anterior part left | 0.003 | 0.004 | 0.004 | 0.448 |
| Gyri parahippocampalis et ambiens anterior part right | 0.010 | 0.004 | 0.010 | 0.042* |
| Superior temporal gyrus   middle part left | 0.001 | 0.002 | 0.001 | 0.541 |
| Superior temporal gyrus middle part right | 0.002 | 0.002 | 0.002 | 0.228 |
| Medial and inferior temporal gyri anterior part left | 0.003 | 0.002 | 0.003 | 0.079 |
| Medial and inferior temporal gyri anterior part right | 0.004 | 0.002 | 0.004 | 0.032* |
| Lateral occipitotemporal gyrus fusiformis anterior part left | 0.006 | 0.005 | 0.007 | 0.231 |
| Lateral occipitotemporal gyrus - gyrus fusiformis anterior part right | 0.008 | 0.004 | 0.008 | 0.125 |
| Cerebellum left | 0.000 | 0.000 | 0.000 | 0.407 |
| Cerebellum right | 0.001 | 0.000 | 0.001 | 0.084 |
| Brainstem | 0.002 | 0.001 | 0.002 | 0.193 |
| Insula right | 0.004 | 0.003 | 0.005 | 0.172 |
| Insula left | 0.006 | 0.003 | 0.006 | 0.073 |
| Occipital lobe right | 0.001 | 0.000 | 0.001 | 0.017* |
| Occipital lobe left | 0.001 | 0.000 | 0.001 | 0.063 |
| Gyri parahippocampalis et ambiens posterior part right | 0.008 | 0.005 | 0.009 | 0.139 |
| Gyri parahippocampalis et ambiens posterior part left | 0.003 | 0.005 | 0.004 | 0.491 |
| Lateral occipitotemporal gyrus - gyrus fusiformis posterior part right | 0.003 | 0.003 | 0.004 | 0.356 |
| Lateral occipitotemporal gyrus fusiformis posterior part left | 0.008 | 0.003 | 0.008 | 0.050* |
| Medial and inferior temporal gyri posterior part right | 0.002 | 0.001 | 0.003 | 0.081 |
| Medial and inferior temporal gyri posterior part left | 0.002 | 0.001 | 0.002 | 0.100 |
| Superior temporal gyrus posterior part right | 0.001 | 0.003 | 0.002 | 0.630 |
| Superior temporal gyrus posterior part left | 0.007 | 0.003 | 0.007 | 0.083 |
| Cingulate gyrus anterior part right | 0.001 | 0.003 | 0.001 | 0.793 |
| Cingulate gyrus anterior part left | 0.002 | 0.002 | 0.002 | 0.376 |
| Cingulate gyrus posterior part right | 0.005 | 0.002 | 0.005 | 0.079 |
| Cingulate gyrus posterior part left | 0.004 | 0.002 | 0.004 | 0.145 |
| Frontal lobe right | 0.000 | 0.000 | 0.000 | 0.094 |
| Frontal lobe left | 0.000 | 0.000 | 0.000 | 0.078 |
| Parietal lobe right | 0.001 | 0.000 | 0.001 | 0.133 |
| Parietal lobe left | 0.001 | 0.000 | 0.001 | 0.105 |
| Caudate nucleus right | 0.002 | 0.002 | 0.002 | 0.541 |
| Caudate nucleus left | 0.002 | 0.002 | 0.002 | 0.539 |
| Thalamus right | 0.003 | 0.002 | 0.003 | 0.126 |
| Thalamus left | 0.004 | 0.002 | 0.004 | 0.035* |
| Subthalamic nucleus right | 0.004 | 0.025 | 0.009 | 0.876 |
| Subthalamic nucleus left | -0.017 | 0.026 | -0.013 | 0.546 |
| Lentiform Nucleus right | 0.002 | 0.002 | 0.002 | 0.382 |
| Lentiform Nucleus left | 0.002 | 0.002 | 0.002 | 0.400 |
| Corpus Callosum | 0.000 | 0.001 | 0.001 | 0.805 |

**Supplementary Table 4. Associations with Bayley-III cognitive subscale after controlling for total brain volume**

Associations between brain volumes at birth and cognitive outcomes at 18-month follow-up after controlling for total brain volume. Linear regression coefficients (β; both standardised and unstandardised), standard errors (SE), and false discovery rate–corrected *p*-values (*p*_FDR_) are reported for each region.

| **Region** | **Unstandardised** β **Coefficient** | **SE** | **Standardised** β **Coefficient** | **p_FDR_** |
| --- | --- | --- | --- | --- |
| Total cortical grey matter | 0.000 | 0.290 | 0.000 | 0.459 |
| Total subcortical grey matter | 0.000 | -0.024 | 0.001 | 0.808 |
| Total white matter | 0.000 | -0.180 | 0.000 | 0.509 |
| Brainstem | -0.001 | -0.025 | 0.002 | 0.807 |
| CSF | 0.000 | -0.020 | 0.000 | 0.807 |
| Ventricles | -0.001 | -0.143 | 0.000 | 0.048* |
| Hippocampus left | 0.007 | 0.072 | 0.007 | 0.417 |
| Hippocampus right | 0.004 | 0.044 | 0.007 | 0.621 |
| Amygdala left | 0.000 | 0.014 | 0.014 | 0.992 |
| Amygdala right | 0.005 | 0.043 | 0.011 | 0.705 |
| Anterior temporal lobe - medial part left | -0.005 | -0.073 | 0.005 | 0.351 |
| Anterior temporal lobe - medial part right | -0.002 | -0.035 | 0.005 | 0.743 |
| Anterior temporal lobe - lateral part left | -0.004 | -0.077 | 0.004 | 0.396 |
| Anterior temporal lobe - lateral part right | -0.005 | -0.085 | 0.005 | 0.410 |
| Gyri parahippocampalis et ambiens anterior part left | -0.002 | -0.019 | 0.005 | 0.727 |
| Gyri parahippocampalis et ambiens anterior part right | 0.007 | 0.102 | 0.005 | 0.244 |
| Superior temporal gyrus -middle part left | -0.002 | -0.095 | 0.002 | 0.396 |
| Superior temporal gyrus- middle part right | 0.000 | -0.013 | 0.002 | 0.901 |
| Medial and inferior temporal gyri anterior part left | 0.002 | 0.079 | 0.002 | 0.456 |
| Medial and inferior temporal gyri anterior part right | 0.003 | 0.144 | 0.002 | 0.240 |
| Lateral occipitotemporal gyrus - gyrus fusiformis anterior part left | 0.001 | 0.013 | 0.006 | 0.886 |
| Lateral occipitotemporal gyrus - gyrus fusiformis anterior part right | 0.004 | 0.055 | 0.005 | 0.534 |
| Cerebellum left | 0.000 | -0.090 | 0.001 | 0.491 |
| Cerebellum right | 0.001 | 0.109 | 0.001 | 0.470 |
| Brainstem | -0.001 | -0.025 | 0.002 | 0.807 |
| Insula right | 0.000 | 0.015 | 0.004 | 0.956 |
| Insula left | 0.003 | 0.097 | 0.004 | 0.455 |
| Occipital lobe right | 0.001 | 0.223 | 0.001 | 0.159 |
| Occipital lobe left | 0.001 | 0.115 | 0.001 | 0.470 |
| Gyri parahippocampalis et ambiens posterior part right | 0.003 | 0.041 | 0.006 | 0.690 |
| Gyri parahippocampalis et ambiens posterior part left | -0.003 | -0.032 | 0.006 | 0.705 |
| Lateral occipitotemporal gyrus - gyrus fusiformis posterior part right | 0.000 | -0.005 | 0.004 | 0.934 |
| Lateral occipitotemporal gyrus- gyrus fusiformis posterior part left | 0.006 | 0.112 | 0.004 | 0.233 |
| Medial and inferior temporal gyri posterior part right | 0.001 | 0.079 | 0.002 | 0.574 |
| Medial and inferior temporal gyri posterior part left | 0.001 | 0.047 | 0.002 | 0.683 |
| Superior temporal gyrus - posterior part right | -0.003 | -0.077 | 0.004 | 0.433 |
| Superior temporal gyrus - posterior part left | 0.004 | 0.088 | 0.004 | 0.352 |
| Cingulate gyrus - anterior part right | -0.005 | -0.133 | 0.003 | 0.205 |
| Cingulate gyrus - anterior part left | -0.001 | -0.025 | 0.003 | 0.775 |
| Cingulate gyrus - posterior part right | 0.002 | 0.065 | 0.003 | 0.514 |
| Cingulate gyrus - posterior part left | 0.001 | 0.029 | 0.003 | 0.886 |
| Frontal lobe right | 0.000 | 0.009 | 0.001 | 0.934 |
| Frontal lobe left | 0.000 | 0.082 | 0.001 | 0.752 |
| Parietal lobe right | 0.000 | -0.075 | 0.001 | 0.814 |
| Parietal lobe left | 0.000 | -0.019 | 0.001 | 0.969 |
| Caudate nucleus right | -0.003 | -0.071 | 0.003 | 0.396 |
| Caudate nucleus left | -0.003 | -0.069 | 0.003 | 0.396 |
| Thalamus right | 0.001 | 0.021 | 0.003 | 0.814 |
| Thalamus left | 0.004 | 0.136 | 0.003 | 0.281 |
| Subthalamic nucleus right | -0.047 | -0.105 | 0.031 | 0.222 |
| Subthalamic nucleus left | -0.064 | -0.127 | 0.031 | 0.145 |
| Lentiform Nucleus right | -0.002 | -0.064 | 0.002 | 0.531 |
| Lentiform Nucleus left | -0.002 | -0.067 | 0.002 | 0.492 |
| Corpus Callosum | -0.003 | -0.109 | 0.002 | 0.230 |

**Supplementary Table 5. Associations with Bayley-III language subscale**

Associations between brain volumes at birth and language outcomes at 18-month follow-up. Linear regression coefficients (β; both standardised and unstandardised), standard errors (SE), and false discovery rate–corrected *p*-values (*p*_FDR_) are reported for each region.

| **Region** | **Estimate** | **SE** | **Std Estimate** | ***p*_FDR_** |
| --- | --- | --- | --- | --- |
| Total brain volume | 0.000 | 0.000 | 0.000 | 0.013* |
| Total white matter | 0.000 | 0.000 | 0.000 | 0.010* |
| Total cortical grey matter | 0.000 | 0.000 | 0.000 | 0.028* |
| Total subcortical grey matter | 0.001 | 0.000 | 0.001 | 0.020* |
| Brainstem | 0.002 | 0.002 | 0.002 | 0.184 |
| Ventricles | 0.000 | 0.001 | 0.000 | 0.895 |
| CSF | 0.000 | 0.000 | 0.000 | 0.136 |
| Hippocampus left | 0.021 | 0.008 | 0.020 | 0.030* |
| Hippocampus right | 0.017 | 0.008 | 0.015 | 0.068 |
| Amygdala left | 0.039 | 0.016 | 0.041 | 0.033* |
| Amygdala right | 0.032 | 0.013 | 0.034 | 0.036* |
| Anterior temporal lobe medial part left | 0.008 | 0.006 | 0.007 | 0.235 |
| Anterior temporal lobe medial part right | 0.010 | 0.006 | 0.010 | 0.093 |
| Anterior temporal lobe lateral part left | 0.008 | 0.005 | 0.008 | 0.184 |
| Anterior temporal lobe lateral part right | 0.006 | 0.005 | 0.006 | 0.317 |
| Gyri parahippocampalis et ambiens anterior part left | 0.014 | 0.006 | 0.014 | 0.031* |
| Gyri parahippocampalis et ambiens anterior part right | 0.015 | 0.006 | 0.015 | 0.030* |
| Superior temporal gyrus middle part left | 0.002 | 0.002 | 0.002 | 0.346 |
| Superior temporal gyrus middle part right | 0.002 | 0.002 | 0.002 | 0.320 |
| Medial and inferior temporal gyri anterior part left | 0.006 | 0.002 | 0.006 | 0.031* |
| Medial and inferior temporal gyri anterior part right | 0.006 | 0.002 | 0.006 | 0.025* |
| Lateral occipitotemporal gyrus fusiformis anterior part left | 0.016 | 0.007 | 0.016 | 0.035* |
| Lateral occipitotemporal gyrus - gyrus fusiformis anterior part right | 0.014 | 0.006 | 0.015 | 0.038* |
| Cerebellum left | 0.001 | 0.001 | 0.001 | 0.130 |
| Cerebellum right | 0.001 | 0.001 | 0.001 | 0.048* |
| Insula right | 0.012 | 0.004 | 0.011 | 0.014* |
| Insula left | 0.011 | 0.004 | 0.011 | 0.011* |
| Occipital lobe right | 0.001 | 0.001 | 0.001 | 0.041* |
| Occipital lobe left | 0.001 | 0.001 | 0.001 | 0.100 |
| Gyri parahippocampalis et ambiens posterior part right | 0.009 | 0.007 | 0.009 | 0.242 |
| Gyri parahippocampalis et ambiens posterior part left | 0.004 | 0.007 | 0.004 | 0.575 |
| Lateral occipitotemporal gyrus   gyrus fusiformis posterior part right | 0.009 | 0.005 | 0.009 | 0.100 |
| Lateral occipitotemporal gyrus fusiformis posterior part left | 0.009 | 0.005 | 0.009 | 0.105 |
| Medial and inferior temporal gyri posterior part right | 0.004 | 0.002 | 0.004 | 0.021* |
| Medial and inferior temporal gyri posterior part left | 0.004 | 0.002 | 0.004 | 0.035* |
| Superior temporal gyrus posterior part right | 0.005 | 0.004 | 0.005 | 0.242 |
| Superior temporal gyrus posterior part left | 0.008 | 0.005 | 0.008 | 0.111 |
| Cingulate gyrus anterior part right | 0.005 | 0.004 | 0.005 | 0.221 |
| Cingulate gyrus anterior part left | 0.003 | 0.003 | 0.003 | 0.468 |
| Cingulate gyrus posterior part right | 0.008 | 0.003 | 0.007 | 0.031* |
| Cingulate gyrus posterior part left | 0.008 | 0.003 | 0.008 | 0.035* |
| Frontal lobe right | 0.001 | 0.000 | 0.001 | 0.064 |
| Frontal lobe left | 0.001 | 0.000 | 0.001 | 0.038* |
| Parietal lobe right | 0.001 | 0.000 | 0.001 | 0.067 |
| Parietal lobe left | 0.001 | 0.000 | 0.001 | 0.019* |
| Caudate nucleus right | 0.010 | 0.003 | 0.010 | 0.011* |
| Caudate nucleus left | 0.009 | 0.003 | 0.009 | 0.025* |
| Thalamus right | 0.005 | 0.002 | 0.006 | 0.049* |
| Thalamus left | 0.006 | 0.002 | 0.006 | 0.028* |
| Subthalamic nucleus right | 0.044 | 0.035 | 0.048 | 0.242 |
| Subthalamic nucleus left | 0.017 | 0.038 | 0.016 | 0.667 |
| Lentiform Nucleus right | 0.005 | 0.003 | 0.005 | 0.057 |
| Lentiform Nucleus left | 0.006 | 0.002 | 0.006 | 0.047* |
| Corpus Callosum | 0.002 | 0.002 | 0.002 | 0.329 |

**Supplementary Table 6. Associations with Bayley-III language subscale after controlling for total brain volume**

Associations between brain volumes at birth and language outcomes at 18-month follow-up after controlling for total brain volume. Linear regression coefficients (β; both standardised and unstandardised), standard errors (SE), and false discovery rate–corrected *p*-values (*p*_FDR_) are reported for each region.

| **Region** | **Unstandardised** β **Coefficient** | **SE** | **Standardised** β **Coefficient** | **p_FDR_** |
| --- | --- | --- | --- | --- |
| Total cortical grey matter | 0.000 | 0.000 | -0.336 | 0.522 |
| Total subcortical grey matter | 0.000 | 0.001 | 0.079 | 0.669 |
| Total white matter | 0.000 | 0.000 | 0.177 | 0.566 |
| Brainstem | -0.003 | 0.003 | -0.127 | 0.342 |
| CSF | 0.000 | 0.000 | 0.031 | 0.769 |
| Ventricles | -0.001 | 0.001 | -0.075 | 0.263 |
| Hippocampus left | 0.011 | 0.010 | 0.063 | 0.424 |
| Hippocampus right | 0.005 | 0.010 | 0.020 | 0.692 |
| Amygdala left | 0.018 | 0.019 | 0.080 | 0.501 |
| Amygdala right | 0.015 | 0.016 | 0.082 | 0.505 |
| Anterior temporal lobe - medial part left | -0.001 | 0.007 | -0.017 | 0.859 |
| Anterior temporal lobe - medial part right | 0.000 | 0.007 | -0.011 | 0.989 |
| Anterior temporal lobe - lateral part left | -0.001 | 0.006 | -0.005 | 0.921 |
| Anterior temporal lobe - lateral part right | -0.005 | 0.007 | -0.069 | 0.540 |
| Gyri parahippocampalis et ambiens anterior part left | 0.007 | 0.007 | 0.077 | 0.421 |
| Gyri parahippocampalis et ambiens anterior part right | 0.007 | 0.007 | 0.071 | 0.470 |
| Superior temporal gyrus -middle part left | -0.004 | 0.003 | -0.122 | 0.368 |
| Superior temporal gyrus- middle part right | -0.004 | 0.003 | -0.125 | 0.342 |
| Medial and inferior temporal gyri anterior part left | 0.002 | 0.003 | 0.070 | 0.572 |
| Medial and inferior temporal gyri anterior part right | 0.002 | 0.003 | 0.093 | 0.540 |
| Lateral occipitotemporal gyrus - gyrus fusiformis anterior part left | 0.008 | 0.008 | 0.075 | 0.477 |
| Lateral occipitotemporal gyrus - gyrus fusiformis anterior part right | 0.006 | 0.007 | 0.073 | 0.533 |
| Cerebellum left | 0.000 | 0.001 | -0.052 | 0.714 |
| Cerebellum right | 0.000 | 0.001 | 0.046 | 0.722 |
| Brainstem | -0.003 | 0.003 | -0.127 | 0.342 |
| Insula right | 0.007 | 0.006 | 0.109 | 0.395 |
| Insula left | 0.007 | 0.005 | 0.114 | 0.349 |
| Occipital lobe right | 0.000 | 0.001 | 0.023 | 0.921 |
| Occipital lobe left | -0.001 | 0.001 | -0.089 | 0.540 |
| Gyri parahippocampalis et ambiens posterior part right | -0.007 | 0.009 | -0.059 | 0.554 |
| Gyri parahippocampalis et ambiens posterior part left | -0.009 | 0.008 | -0.070 | 0.412 |
| Lateral occipitotemporal gyrus - gyrus fusiformis posterior part right | 0.002 | 0.006 | 0.036 | 0.721 |
| Lateral occipitotemporal gyrus- gyrus fusiformis posterior part left | 0.002 | 0.006 | 0.036 | 0.714 |
| Medial and inferior temporal gyri posterior part right | 0.002 | 0.003 | 0.087 | 0.572 |
| Medial and inferior temporal gyri posterior part left | 0.001 | 0.002 | 0.023 | 0.762 |
| Superior temporal gyrus - posterior part right | -0.003 | 0.005 | -0.055 | 0.645 |
| Superior temporal gyrus - posterior part left | 0.001 | 0.005 | 0.013 | 0.811 |
| Cingulate gyrus - anterior part right | -0.004 | 0.005 | -0.072 | 0.528 |
| Cingulate gyrus - anterior part left | -0.005 | 0.004 | -0.068 | 0.404 |
| Cingulate gyrus - posterior part right | 0.002 | 0.005 | 0.035 | 0.670 |
| Cingulate gyrus - posterior part left | 0.002 | 0.005 | 0.039 | 0.709 |
| Frontal lobe right | -0.001 | 0.001 | -0.357 | 0.222 |
| Frontal lobe left | -0.001 | 0.001 | -0.208 | 0.540 |
| Parietal lobe right | -0.001 | 0.001 | -0.249 | 0.395 |
| Parietal lobe left | 0.000 | 0.001 | 0.041 | 0.747 |
| Caudate nucleus right | 0.006 | 0.004 | 0.112 | 0.282 |
| Caudate nucleus left | 0.004 | 0.004 | 0.074 | 0.495 |
| Thalamus right | 0.000 | 0.004 | 0.008 | 0.992 |
| Thalamus left | 0.002 | 0.004 | 0.065 | 0.666 |
| Subthalamic nucleus right | -0.031 | 0.044 | -0.042 | 0.572 |
| Subthalamic nucleus left | -0.056 | 0.044 | -0.082 | 0.342 |
| Lentiform Nucleus right | 0.001 | 0.003 | 0.024 | 0.837 |
| Lentiform Nucleus left | 0.001 | 0.003 | 0.034 | 0.751 |
| Corpus Callosum | -0.003 | 0.002 | -0.074 | 0.424 |

**Supplementary Table 7. Associations with Bayley-III motor subscale**

Associations between brain volumes at birth and motor outcomes at 18-month follow-up. Linear regression coefficients (β; both standardised and unstandardised), standard errors (SE), and false discovery rate–corrected *p*-values (*p*_FDR_) are reported for each region.

| **Region** | **Unstandardised** β **Coefficient** | **SE** | **Standardised** β **Coefficient** | **p_FDR_** |
| --- | --- | --- | --- | --- |
| Total brain volume | 0.000 | 0.000 | 0.000 | 0.846 |
| Total white matter volume | 0.000 | 0.000 | 0.000 | 0.821 |
| Total cortical grey matter | 0.000 | 0.000 | 0.000 | 0.718 |
| Total subcortical grey matter | 0.000 | 0.000 | 0.000 | 0.884 |
| Brainstem | -0.001 | 0.001 | -0.001 | 0.417 |
| Ventricles | -0.001 | 0.000 | -0.001 | 0.023* |
| CSF | 0.000 | 0.000 | 0.000 | 0.779 |
| Hippocampus left | 0.002 | 0.005 | 0.002 | 0.884 |
| Hippocampus right | 0.004 | 0.005 | 0.004 | 0.732 |
| Amygdala left | 0.011 | 0.010 | 0.014 | 0.476 |
| Amygdala right | 0.002 | 0.008 | 0.004 | 0.885 |
| Anterior temporal lobe medial part left | 0.002 | 0.004 | 0.003 | 0.838 |
| Anterior temporal lobe medial part right | 0.005 | 0.003 | 0.006 | 0.291 |
| Anterior temporal lobe lateral part left | 0.002 | 0.003 | 0.002 | 0.838 |
| Anterior temporal lobe lateral part right | 0.000 | 0.003 | 0.001 | 0.944 |
| Gyri parahippocampalis et ambiens anterior part left | 0.001 | 0.004 | 0.002 | 0.884 |
| Gyri parahippocampalis et ambiens anterior part right | 0.000 | 0.004 | 0.000 | 0.978 |
| Superior temporal gyrus middle part left | -0.001 | 0.001 | -0.001 | 0.732 |
| Superior temporal gyrus middle part right | 0.000 | 0.001 | 0.000 | 0.948 |
| Medial and inferior temporal gyri anterior part left | 0.001 | 0.001 | 0.001 | 0.826 |
| Medial and inferior temporal gyri anterior part right | 0.002 | 0.001 | 0.002 | 0.291 |
| Lateral occipitotemporal gyrus fusiformis anterior part left | -0.001 | 0.004 | -0.001 | 0.890 |
| Lateral occipitotemporal gyrus - gyrus fusiformis anterior part right | 0.003 | 0.004 | 0.003 | 0.732 |
| Cerebellum left | -0.001 | 0.000 | -0.001 | 0.230 |
| Cerebellum right | 0.000 | 0.000 | 0.000 | 0.573 |
| Brainstem | -0.001 | 0.001 | -0.001 | 0.417 |
| Insula right | 0.002 | 0.003 | 0.002 | 0.679 |
| Insula left | 0.002 | 0.002 | 0.002 | 0.681 |
| Occipital lobe right | 0.000 | 0.000 | 0.000 | 0.417 |
| Occipital lobe left | 0.000 | 0.000 | 0.000 | 0.476 |
| Gyri parahippocampalis et ambiens posterior part right | 0.001 | 0.004 | 0.001 | 0.917 |
| Gyri parahippocampalis et ambiens posterior part left | 0.000 | 0.004 | 0.000 | 0.981 |
| Lateral occipitotemporal gyrus   gyrus fusiformis posterior part right | -0.001 | 0.003 | -0.001 | 0.884 |
| Lateral occipitotemporal gyrus fusiformis posterior part left | 0.003 | 0.003 | 0.003 | 0.653 |
| Medial and inferior temporal gyri posterior part right | 0.001 | 0.001 | 0.001 | 0.494 |
| Medial and inferior temporal gyri posterior part left | 0.001 | 0.001 | 0.001 | 0.594 |
| Superior temporal gyrus posterior part right | -0.001 | 0.003 | -0.001 | 0.884 |
| Superior temporal gyrus posterior part left | 0.002 | 0.003 | 0.002 | 0.820 |
| Cingulate gyrus anterior part right | -0.001 | 0.002 | -0.001 | 0.884 |
| Cingulate gyrus anterior part left | 0.000 | 0.002 | 0.000 | 0.942 |
| Cingulate gyrus posterior part right | 0.003 | 0.002 | 0.003 | 0.278 |
| Cingulate gyrus posterior part left | 0.003 | 0.002 | 0.003 | 0.294 |
| Frontal lobe right | 0.000 | 0.000 | 0.000 | 0.884 |
| Frontal lobe left | 0.000 | 0.000 | 0.000 | 0.884 |
| Parietal lobe right | 0.000 | 0.000 | 0.000 | 0.855 |
| Parietal lobe left | 0.000 | 0.000 | 0.000 | 0.658 |
| Caudate nucleus right | 0.000 | 0.002 | 0.000 | 0.960 |
| Caudate nucleus left | 0.000 | 0.002 | 0.001 | 0.917 |
| Thalamus right | 0.002 | 0.002 | 0.002 | 0.567 |
| Thalamus left | 0.002 | 0.002 | 0.002 | 0.494 |
| Subthalamic nucleus right | 0.022 | 0.022 | 0.027 | 0.567 |
| Subthalamic nucleus left | 0.009 | 0.024 | 0.013 | 0.884 |
| Lentiform Nucleus right | 0.001 | 0.002 | 0.001 | 0.884 |
| Lentiform Nucleus left | 0.000 | 0.002 | 0.000 | 0.924 |
| Corpus Callosum | -0.002 | 0.001 | -0.001 | 0.384 |

**Supplementary Table 8. Associations with Bayley-III motor subscale after controlling for total brain volume**

Associations between brain volumes at birth and language outcomes at 18-month follow-up after controlling for total brain volume. Linear regression coefficients (β; both standardised and unstandardised), standard errors (SE), and false discovery rate–corrected *p*-values (*p*_FDR_) are reported for each region.

| **Region** | **Unstandardised** β **Coefficient** | **SE** | **Standardised** β **Coefficient** | **p_FDR_** |
| --- | --- | --- | --- | --- |
| Brainstem | -0.004 | 0.002 | -0.286 | 0.039* |
| CSF | 0.000 | 0.000 | -0.070 | 0.626 |
| Cortical gray matter | 0.000 | 0.000 | 0.347 | 0.647 |
| Deep gray matter | 0.000 | 0.000 | -0.013 | 0.939 |
| Ventricles | -0.001 | 0.000 | -0.184 | 0.006** |
| White matter | 0.000 | 0.000 | 0.082 | 0.939 |
| Hippocampus left | 0.000 | 0.006 | 0.004 | 0.983 |
| Hippocampus right | 0.003 | 0.006 | 0.037 | 0.939 |
| Amygdala left | 0.013 | 0.012 | 0.091 | 0.626 |
| Amygdala right | 0.000 | 0.010 | 0.007 | 0.988 |
| Anterior temporal lobe - medial part left | 0.002 | 0.004 | 0.033 | 0.939 |
| Anterior temporal lobe - medial part right | 0.007 | 0.005 | 0.132 | 0.347 |
| Anterior temporal lobe - lateral part left | 0.001 | 0.004 | 0.031 | 0.939 |
| Anterior temporal lobe - lateral part right | -0.001 | 0.004 | -0.024 | 0.939 |
| Gyri parahippocampalis et ambiens anterior part left | 0.000 | 0.004 | 0.010 | 0.983 |
| Gyri parahippocampalis et ambiens anterior part right | -0.002 | 0.005 | -0.028 | 0.939 |
| Superior temporal gyrus -middle part left | -0.003 | 0.002 | -0.146 | 0.363 |
| Superior temporal gyrus- middle part right | -0.001 | 0.002 | -0.038 | 0.939 |
| Medial and inferior temporal gyri anterior part left | 0.001 | 0.002 | 0.027 | 0.939 |
| Medial and inferior temporal gyri anterior part right | 0.003 | 0.002 | 0.173 | 0.319 |
| Lateral occipitotemporal gyrus - gyrus fusiformis anterior part left | -0.004 | 0.005 | -0.055 | 0.850 |
| Lateral occipitotemporal gyrus - gyrus fusiformis anterior part right | 0.002 | 0.005 | 0.042 | 0.939 |
| Cerebellum left | -0.002 | 0.001 | -0.348 | 0.025* |
| Cerebellum right | -0.001 | 0.001 | -0.218 | 0.263 |
| Brainstem | -0.004 | 0.002 | -0.286 | 0.039* |
| Insula right | 0.002 | 0.004 | 0.071 | 0.891 |
| Insula left | 0.002 | 0.003 | 0.067 | 0.906 |
| Occipital lobe right | 0.001 | 0.001 | 0.171 | 0.433 |
| Occipital lobe left | 0.001 | 0.001 | 0.165 | 0.505 |
| Gyri parahippocampalis et ambiens posterior part right | -0.001 | 0.006 | -0.011 | 0.944 |
| Gyri parahippocampalis et ambiens posterior part left | -0.002 | 0.005 | -0.025 | 0.939 |
| Lateral occipitotemporal gyrus - gyrus fusiformis posterior part right | -0.002 | 0.004 | -0.053 | 0.858 |
| Lateral occipitotemporal gyrus - gyrus fusiformis posterior part left | 0.002 | 0.003 | 0.058 | 0.849 |
| Medial and inferior temporal gyri posterior part right | 0.002 | 0.002 | 0.144 | 0.556 |
| Medial and inferior temporal gyri posterior part left | 0.001 | 0.002 | 0.092 | 0.742 |
| Superior temporal gyrus - posterior part right | -0.003 | 0.003 | -0.075 | 0.726 |
| Superior temporal gyrus - posterior part left | 0.001 | 0.003 | 0.030 | 0.939 |
| Cingulate gyrus - anterior part right | -0.003 | 0.003 | -0.079 | 0.710 |
| Cingulate gyrus- anterior part left | -0.001 | 0.003 | -0.036 | 0.939 |
| Cingulate gyrus- posterior part right | 0.005 | 0.003 | 0.159 | 0.305 |
| Cingulate gyrus- posterior part left | 0.005 | 0.003 | 0.171 | 0.319 |
| Frontal lobe right | 0.000 | 0.000 | -0.057 | 0.939 |
| Frontal lobe left | 0.000 | 0.001 | -0.062 | 0.939 |
| Parietal lobe right | 0.000 | 0.001 | -0.007 | 0.968 |
| Parietal lobe left | 0.001 | 0.001 | 0.161 | 0.713 |
| Caudate nucleus right | -0.002 | 0.003 | -0.038 | 0.932 |
| Caudate nucleus left | 0.000 | 0.003 | -0.006 | 0.939 |
| Thalamus right | 0.002 | 0.002 | 0.099 | 0.698 |
| Thalamus left | 0.003 | 0.002 | 0.122 | 0.576 |
| Subthalamic nucleus right | 0.024 | 0.028 | 0.081 | 0.741 |
| Subthalamic nucleus left | 0.003 | 0.028 | 0.017 | 0.944 |
| Lentiform Nucleus right | 0.000 | 0.002 | 0.002 | 0.978 |
| Lentiform Nucleus left | -0.002 | 0.002 | -0.069 | 0.789 |
| Corpus Callosum | -0.003 | 0.002 | -0.169 | 0.123 |

**Supplementary Table 9. Associations with internalising**

Associations between brain volumes at birth and internalising at 18-month follow-up. Linear regression coefficients (β; both standardised and unstandardised), standard errors (SE), and false discovery rate–corrected *p*-values (*p*_FDR_) are reported for each region.

| **Region** | **Unstandardised** β **Coefficient** | **SE** | **Standardised** β **Coefficient** | **p_FDR_** |
| --- | --- | --- | --- | --- |
| Total | 0.000 | 0.000 | 0.000 | 0.463 |
| ICV | 0.000 | 0.000 | 0.000 | 0.513 |
| White matter | 0.000 | 0.000 | 0.000 | 0.474 |
| Deep Gray Matter | 0.000 | 0.000 | 0.000 | 0.445 |
| Brainstem | -0.001 | 0.001 | -0.001 | 0.426 |
| Ventricles | 0.000 | 0.000 | 0.000 | 0.663 |
| CSF | 0.000 | 0.000 | 0.000 | 0.998 |
| Cortical gray matter | 0.000 | 0.000 | 0.000 | 0.513 |
| Hippocampus left | -0.004 | 0.003 | -0.004 | 0.470 |
| Hippocampus right | -0.004 | 0.003 | -0.004 | 0.463 |
| Amygdala left | -0.007 | 0.005 | -0.005 | 0.508 |
| Amygdala right | -0.010 | 0.004 | -0.008 | 0.241 |
| Anterior temporal lobe medial part left | -0.001 | 0.002 | 0.000 | 0.888 |
| Anterior temporal lobe medial part right | 0.000 | 0.002 | 0.000 | 0.945 |
| Anterior temporal lobe lateral part left | 0.000 | 0.002 | 0.001 | 0.970 |
| Anterior temporal lobe lateral part right | -0.001 | 0.002 | -0.001 | 0.865 |
| Gyri parahippocampalis et ambiens anterior part left | -0.003 | 0.002 | -0.003 | 0.380 |
| Gyri parahippocampalis et ambiens anterior part right | -0.005 | 0.002 | -0.004 | 0.205 |
| Superior temporal gyrus middle part left | 0.000 | 0.001 | 0.000 | 0.922 |
| Superior temporal gyrus middle part right | -0.001 | 0.001 | -0.001 | 0.610 |
| Medial and inferior temporal gyri anterior part left | -0.001 | 0.001 | -0.001 | 0.495 |
| Medial and inferior temporal gyri anterior part right | 0.000 | 0.001 | 0.000 | 0.769 |
| Lateral occipitotemporal gyrus fusiformis anterior part left | -0.002 | 0.002 | -0.001 | 0.665 |
| Lateral occipitotemporal gyrus   gyrus fusiformis anterior part right | -0.001 | 0.002 | -0.001 | 0.792 |
| Cerebellum left | 0.000 | 0.000 | 0.000 | 0.241 |
| Cerebellum right | 0.000 | 0.000 | 0.000 | 0.241 |
| Brainstem | -0.001 | 0.001 | -0.001 | 0.426 |
| Insula right | -0.002 | 0.001 | -0.002 | 0.432 |
| Insula left | -0.001 | 0.001 | -0.001 | 0.560 |
| Occipital lobe right | 0.000 | 0.000 | 0.000 | 0.430 |
| Occipital lobe left | 0.000 | 0.000 | 0.000 | 0.455 |
| Gyri parahippocampalis et ambiens posterior part right | -0.004 | 0.002 | -0.003 | 0.425 |
| Gyri parahippocampalis et ambiens posterior part left | 0.000 | 0.002 | 0.000 | 0.982 |
| Lateral occipitotemporal gyrus   gyrus fusiformis posterior part right | -0.002 | 0.002 | -0.002 | 0.513 |
| Lateral occipitotemporal gyrus fusiformis posterior part left | -0.003 | 0.002 | -0.003 | 0.317 |
| Medial and inferior temporal gyri posterior part right | 0.000 | 0.001 | 0.000 | 0.576 |
| Medial and inferior temporal gyri posterior part left | 0.000 | 0.001 | 0.000 | 0.823 |
| Superior temporal gyrus posterior part right | -0.001 | 0.001 | -0.001 | 0.720 |
| Superior temporal gyrus posterior part left | -0.002 | 0.002 | -0.002 | 0.426 |
| Cingulate gyrus anterior part right | 0.000 | 0.001 | 0.000 | 0.899 |
| Cingulate gyrus anterior part left | 0.000 | 0.001 | 0.000 | 0.899 |
| Cingulate gyrus posterior part right | -0.001 | 0.001 | -0.001 | 0.513 |
| Cingulate gyrus posterior part left | -0.002 | 0.001 | -0.001 | 0.450 |
| Frontal lobe right | 0.000 | 0.000 | 0.000 | 0.865 |
| Frontal lobe left | 0.000 | 0.000 | 0.000 | 0.820 |
| Parietal lobe right | 0.000 | 0.000 | 0.000 | 0.513 |
| Parietal lobe left | 0.000 | 0.000 | 0.000 | 0.459 |
| Caudate nucleus right | -0.001 | 0.001 | -0.001 | 0.513 |
| Caudate nucleus left | -0.001 | 0.001 | -0.001 | 0.513 |
| Thalamus right | -0.001 | 0.001 | -0.001 | 0.380 |
| Thalamus left | -0.002 | 0.001 | -0.001 | 0.329 |
| Subthalamic nucleus right | 0.000 | 0.012 | 0.004 | 0.982 |
| Subthalamic nucleus left | 0.002 | 0.012 | 0.006 | 0.945 |
| Lentiform Nucleus right | -0.001 | 0.001 | 0.000 | 0.638 |
| Lentiform Nucleus left | -0.001 | 0.001 | -0.001 | 0.575 |
| Corpus Callosum | 0.000 | 0.001 | 0.000 | 0.984 |

**Supplementary Table 10. Associations with internalising after controlling for total brain volume**

Associations between brain volumes at birth and internalising at 18-month follow-up after controlling for total brain volume. Linear regression coefficients (β; both standardised and unstandardised), standard errors (SE), and false discovery rate–corrected *p*-values (*p*_FDR_) are reported for each region.

| **Region** | **Unstandardised** β **Coefficient** | **SE** | **Standardised** β **Coefficient** | **p_FDR_** |
| --- | --- | --- | --- | --- |
| Brainstem | -0.001 | 0.001 | -0.079 | 0.741 |
| CSF | 0.000 | 0.000 | 0.054 | 0.709 |
| Cortical gray matter | 0.000 | 0.000 | 0.408 | 0.583 |
| Deep gray matter | 0.000 | 0.000 | -0.060 | 0.874 |
| Ventricles | 0.000 | 0.000 | 0.089 | 0.583 |
| White matter | 0.000 | 0.000 | -0.011 | 0.966 |
| Cerebellum | 0.000 | 0.000 | -0.219 | 0.556 |
| Hippocampus left | -0.003 | 0.003 | -0.056 | 0.709 |
| Hippocampus right | -0.003 | 0.003 | -0.060 | 0.709 |
| Amygdala left | -0.002 | 0.006 | -0.024 | 0.874 |
| Amygdala right | -0.008 | 0.005 | -0.112 | 0.583 |
| Anterior temporal lobe - medial part left | 0.001 | 0.002 | 0.042 | 0.847 |
| Anterior temporal lobe - medial part right | 0.002 | 0.002 | 0.080 | 0.699 |
| Anterior temporal lobe - lateral part left | 0.002 | 0.002 | 0.094 | 0.583 |
| Anterior temporal lobe - lateral part right | 0.001 | 0.002 | 0.047 | 0.856 |
| Gyri parahippocampalis et ambiens anterior part left | -0.002 | 0.002 | -0.061 | 0.699 |
| Gyri parahippocampalis et ambiens anterior part right | -0.004 | 0.002 | -0.133 | 0.556 |
| Superior temporal gyrus -middle part left | 0.002 | 0.001 | 0.165 | 0.583 |
| Superior temporal gyrus- middle part right | 0.000 | 0.001 | 0.024 | 0.874 |
| Medial and inferior temporal gyri anterior part left | -0.001 | 0.001 | -0.056 | 0.847 |
| Medial and inferior temporal gyri anterior part right | 0.001 | 0.001 | 0.060 | 0.851 |
| Lateral occipitotemporal gyrus - gyrus fusiformis anterior part left | 0.000 | 0.003 | 0.001 | 0.990 |
| Lateral occipitotemporal gyrus - gyrus fusiformis anterior part right | 0.001 | 0.002 | 0.030 | 0.874 |
| Cerebellum left | 0.000 | 0.000 | -0.210 | 0.563 |
| Cerebellum right | 0.000 | 0.000 | -0.210 | 0.556 |
| Brainstem | -0.001 | 0.001 | -0.079 | 0.741 |
| Insula right | -0.001 | 0.002 | -0.077 | 0.739 |
| Insula left | 0.000 | 0.002 | 0.021 | 0.875 |
| Occipital lobe right | 0.000 | 0.000 | -0.068 | 0.847 |
| Occipital lobe left GM | 0.000 | 0.000 | -0.064 | 0.874 |
| Gyri parahippocampalis et ambiens posterior part right | -0.003 | 0.003 | -0.079 | 0.679 |
| Gyri parahippocampalis et ambiens posterior part left | 0.002 | 0.003 | 0.057 | 0.709 |
| Lateral occipitotemporal gyrus- gyrus fusiformis posterior part right | -0.001 | 0.002 | -0.029 | 0.874 |
| Lateral occipitotemporal gyrus- gyrus fusiformis posterior part left | -0.002 | 0.002 | -0.100 | 0.583 |
| Medial and inferior temporal gyri posterior part right | 0.000 | 0.001 | 0.044 | 0.874 |
| Medial and inferior temporal gyri posterior part left | 0.001 | 0.001 | 0.101 | 0.709 |
| Superior temporal gyrus - posterior part right | 0.000 | 0.002 | 0.020 | 0.874 |
| Superior temporal gyrus - posterior part left | -0.002 | 0.002 | -0.090 | 0.614 |
| Cingulate gyrus- anterior part right | 0.003 | 0.002 | 0.130 | 0.574 |
| Cingulate gyrus- anterior part left | 0.002 | 0.001 | 0.091 | 0.583 |
| Cingulate gyrus- posterior part right | 0.000 | 0.002 | -0.026 | 0.874 |
| Cingulate gyrus- posterior part left | -0.001 | 0.002 | -0.043 | 0.874 |
| Frontal lobe right | 0.001 | 0.000 | 0.528 | 0.556 |
| Frontal lobe left | 0.001 | 0.000 | 0.485 | 0.556 |
| Parietal lobe right | 0.000 | 0.000 | 0.028 | 0.911 |
| Parietal lobe left | 0.000 | 0.000 | -0.109 | 0.856 |
| Caudate nucleus right | 0.000 | 0.001 | -0.025 | 0.874 |
| Caudate nucleus left | 0.000 | 0.001 | -0.017 | 0.875 |
| Thalamus right | -0.001 | 0.001 | -0.115 | 0.643 |
| Thalamus left | -0.002 | 0.001 | -0.148 | 0.583 |
| Subthalamic nucleus right | 0.021 | 0.015 | 0.113 | 0.583 |
| Subthalamic nucleus left | 0.020 | 0.015 | 0.091 | 0.583 |
| Lentiform Nucleus right | 0.000 | 0.001 | 0.036 | 0.874 |
| Lentiform Nucleus left | 0.000 | 0.001 | 0.022 | 0.874 |
| Corpus Callosum | 0.001 | 0.001 | 0.095 | 0.583 |

**Supplementary Table 11. Associations with externalising**

Associations between brain volumes at birth and externalising at 18-month follow-up. Linear regression coefficients (β; both standardised and unstandardised), standard errors (SE), and false discovery rate–corrected *p*-values (*p*_FDR_) are reported for each region.

| **Region** | **Unstandardised** β **Coefficient** | **SE** | **Standardised** β **Coefficient** | **p_FDR_** |
| --- | --- | --- | --- | --- |
| Total | 0.000 | 0.000 | 0.000 | 0.982 |
| ICV | 0.000 | 0.000 | 0.000 | 0.984 |
| White matter | 0.000 | 0.000 | 0.000 | 0.982 |
| Deep Gray Matter | 0.000 | 0.000 | 0.000 | 0.984 |
| Brainstem | 0.000 | 0.001 | 0.000 | 0.946 |
| Ventricles | 0.000 | 0.000 | 0.000 | 0.837 |
| CSF | 0.000 | 0.000 | 0.000 | 0.994 |
| Cortical gray matter | 0.000 | 0.000 | 0.000 | 0.984 |
| Hippocampus left | -0.001 | 0.004 | -0.001 | 0.982 |
| Hippocampus right | -0.001 | 0.004 | -0.001 | 0.984 |
| Amygdala left | -0.005 | 0.007 | -0.001 | 0.837 |
| Amygdala right | -0.006 | 0.006 | -0.004 | 0.837 |
| Anterior temporal lobe medial part left | 0.003 | 0.003 | 0.004 | 0.837 |
| Anterior temporal lobe medial part right | 0.005 | 0.003 | 0.005 | 0.367 |
| Anterior temporal lobe lateral part left | -0.002 | 0.002 | -0.002 | 0.837 |
| Anterior temporal lobe lateral part right | -0.001 | 0.003 | -0.001 | 0.915 |
| Gyri parahippocampalis et ambiens anterior part left | 0.001 | 0.003 | 0.002 | 0.946 |
| Gyri parahippocampalis et ambiens anterior part right | 0.000 | 0.003 | 0.001 | 0.985 |
| Superior temporal gyrus middle part left | 0.000 | 0.001 | 0.000 | 0.985 |
| Superior temporal gyrus middle part right | 0.000 | 0.001 | 0.001 | 0.946 |
| Medial and inferior temporal gyri anterior part left | -0.001 | 0.001 | -0.001 | 0.837 |
| Medial and inferior temporal gyri anterior part right | 0.001 | 0.001 | 0.001 | 0.837 |
| Lateral occipitotemporal gyrus fusiformis anterior part left | 0.000 | 0.003 | 0.000 | 0.985 |
| Lateral occipitotemporal gyrus   gyrus fusiformis anterior part right | -0.001 | 0.003 | 0.000 | 0.976 |
| Cerebellum left | 0.000 | 0.000 | 0.000 | 0.915 |
| Cerebellum right | 0.000 | 0.000 | 0.000 | 0.965 |
| Brainstem | 0.000 | 0.001 | 0.000 | 0.946 |
| Insula right | -0.001 | 0.002 | -0.001 | 0.930 |
| Insula left | 0.000 | 0.002 | 0.001 | 0.984 |
| Occipital lobe right | 0.000 | 0.000 | 0.000 | 0.837 |
| Occipital lobe left | 0.000 | 0.000 | 0.000 | 0.984 |
| Gyri parahippocampalis et ambiens posterior part right | -0.001 | 0.003 | 0.000 | 0.982 |
| Gyri parahippocampalis et ambiens posterior part left | 0.000 | 0.003 | 0.000 | 0.984 |
| Lateral occipitotemporal gyrus – gyrus fusiformis posterior part right | 0.001 | 0.002 | 0.001 | 0.867 |
| Lateral occipitotemporal gyrus fusiformis posterior part left | 0.001 | 0.002 | 0.001 | 0.954 |
| Medial and inferior temporal gyri posterior part | 0.000 | 0.001 | 0.000 | 0.910 |
| Medial and inferior temporal gyri posterior part left | 0.000 | 0.001 | 0.000 | 0.984 |
| Superior temporal gyrus posterior part right | 0.000 | 0.002 | 0.001 | 0.982 |
| Superior temporal gyrus posterior part left | 0.000 | 0.002 | 0.000 | 0.985 |
| Cingulate gyrus anterior part right | 0.000 | 0.002 | 0.000 | 0.985 |
| Cingulate gyrus anterior part left | 0.000 | 0.002 | 0.000 | 0.984 |
| Cingulate gyrus posterior part right | -0.001 | 0.001 | -0.001 | 0.837 |
| Cingulate gyrus posterior part left | 0.000 | 0.002 | 0.001 | 0.985 |
| Frontal lobe right | 0.000 | 0.000 | 0.000 | 0.985 |
| Frontal lobe left | 0.000 | 0.000 | 0.000 | 0.985 |
| Parietal lobe right | 0.000 | 0.000 | 0.000 | 0.985 |
| Parietal lobe left | 0.000 | 0.000 | 0.000 | 0.837 |
| Caudate nucleus right | 0.001 | 0.002 | 0.001 | 0.837 |
| Caudate nucleus left | 0.000 | 0.002 | 0.001 | 0.982 |
| Thalamus right | 0.000 | 0.001 | 0.000 | 0.945 |
| Thalamus left | 0.000 | 0.001 | 0.000 | 0.987 |
| Subthalamic nucleus right | 0.000 | 0.016 | 0.009 | 0.990 |
| Subthalamic nucleus left | -0.001 | 0.018 | 0.006 | 0.985 |
| Lentiform Nucleus right | 0.001 | 0.001 | 0.001 | 0.837 |
| Lentiform Nucleus left | 0.001 | 0.001 | 0.001 | 0.837 |
| Corpus Callosum | 0.001 | 0.001 | 0.001 | 0.842 |

**Supplementary Table 12. Associations with externalising after controlling for total brain volume**

Associations between brain volumes at birth and externalising at 18-month follow-up after controlling for total brain volume. Linear regression coefficients (β; both standardised and unstandardised), standard errors (SE), and false discovery rate–corrected *p*-values (*p*_FDR_) are reported for each region.

| **Region** | **Unstandardised** β **Coefficient** | **SE** | **Standardised** β **Coefficient** | **p_FDR_** |
| --- | --- | --- | --- | --- |
| Brainstem | 0.001 | 0.001 | 0.095 | 0.912 |
| CSF | 0.000 | 0.000 | 0.007 | 0.986 |
| Total white matter | 0.000 | 0.000 | -0.009 | 0.999 |
| Total cortical grey matter | 0.000 | 0.000 | -0.002 | 1.000 |
| Total subcortical grey matter | 0.000 | 0.000 | 0.108 | 0.913 |
| Ventricles | 0.000 | 0.000 | 0.063 | 0.913 |
| Hippocampus left | -0.002 | 0.005 | -0.024 | 0.974 |
| Hippocampus right | -0.001 | 0.005 | -0.020 | 0.974 |
| Amygdala left | -0.002 | 0.009 | -0.019 | 0.974 |
| Amygdala right | -0.006 | 0.008 | -0.059 | 0.913 |
| Anterior temporal lobe - medial part left | 0.005 | 0.003 | 0.115 | 0.679 |
| Anterior temporal lobe - medial part right | 0.008 | 0.003 | 0.209 | 0.505 |
| Anterior temporal lobe - lateral part left | -0.003 | 0.003 | -0.078 | 0.913 |
| Anterior temporal lobe - lateral part right | -0.001 | 0.003 | -0.039 | 0.974 |
| Gyri parahippocampalis et ambiens anterior part left | 0.003 | 0.003 | 0.061 | 0.913 |
| Gyri parahippocampalis et ambiens anterior part right | 0.002 | 0.003 | 0.034 | 0.974 |
| Superior temporal gyrus – middle part left | 0.000 | 0.001 | 0.022 | 0.976 |
| Superior temporal gyrus – middle part right | 0.001 | 0.001 | 0.073 | 0.913 |
| Medial and inferior temporal gyri anterior part left | -0.001 | 0.001 | -0.105 | 0.913 |
| Medial and inferior temporal gyri anterior part right | 0.003 | 0.001 | 0.208 | 0.505 |
| Lateral occipitotemporal gyrus – gyrus fusiformis anterior part left | 0.000 | 0.004 | 0.007 | 0.990 |
| Lateral occipitotemporal gyrus – gyrus fusiformis anterior part right | 0.000 | 0.003 | -0.004 | 0.990 |
| Cerebellum left | 0.000 | 0.000 | -0.067 | 0.959 |
| Cerebellum right | 0.000 | 0.000 | -0.049 | 0.974 |
| Insula right | -0.001 | 0.003 | -0.056 | 0.958 |
| Insula left | 0.001 | 0.002 | 0.037 | 0.974 |
| Occipital lobe right | -0.001 | 0.000 | -0.179 | 0.726 |
| Occipital lobe left | 0.000 | 0.000 | 0.005 | 0.999 |
| Gyri parahippocampalis et ambiens posterior part right | -0.001 | 0.004 | -0.019 | 0.974 |
| Gyri parahippocampalis et ambiens posterior part left | 0.001 | 0.004 | 0.011 | 0.976 |
| Lateral occipitotemporal gyrus – gyrus fusiformis posterior part right | 0.002 | 0.003 | 0.046 | 0.952 |
| Lateral occipitotemporal gyrus – gyrus fusiformis posterior part left | 0.001 | 0.003 | 0.032 | 0.974 |
| Medial and inferior temporal gyri posterior part right | -0.001 | 0.001 | -0.079 | 0.932 |
| Medial and inferior temporal gyri posterior part left | 0.000 | 0.001 | -0.003 | 1.000 |
| Superior temporal gyrus – posterior part right | 0.001 | 0.002 | 0.044 | 0.960 |
| Superior temporal gyrus – posterior part left | 0.000 | 0.003 | -0.002 | 1.000 |
| Cingulate gyrus – anterior part right | 0.000 | 0.002 | 0.008 | 0.988 |
| Cingulate gyrus – anterior part left | 0.001 | 0.002 | 0.024 | 0.974 |
| Cingulate gyrus – posterior part right | -0.002 | 0.002 | -0.080 | 0.913 |
| Cingulate gyrus – posterior part left | 0.001 | 0.002 | 0.064 | 0.925 |
| Frontal lobe right | 0.000 | 0.000 | 0.138 | 0.932 |
| Frontal lobe left | 0.000 | 0.000 | 0.138 | 0.932 |
| Parietal lobe right | 0.000 | 0.000 | 0.028 | 0.976 |
| Parietal lobe left | -0.001 | 0.000 | -0.255 | 0.913 |
| Caudate nucleus right | 0.002 | 0.002 | 0.093 | 0.913 |
| Caudate nucleus left | 0.001 | 0.002 | 0.056 | 0.913 |
| Thalamus right | -0.001 | 0.002 | -0.052 | 0.974 |
| Thalamus left | 0.000 | 0.002 | 0.026 | 0.974 |
| Subthalamic nucleus right | 0.013 | 0.021 | 0.050 | 0.928 |
| Subthalamic nucleus left | 0.008 | 0.021 | 0.025 | 0.974 |
| Lentiform Nucleus right | 0.002 | 0.002 | 0.151 | 0.700 |
| Lentiform Nucleus left | 0.003 | 0.002 | 0.162 | 0.535 |
| Corpus Callosum | 0.001 | 0.001 | 0.072 | 0.913 |

**Supplementary Table 13. Brain volumes mediating the association between sex and language outcomes**

Associations between brain volumes at birth and internalising at 18-month follow-up after controlling for total brain volume. Linear regression coefficients (β; both standardised and unstandardised), standard errors (SE), and false discovery rate–corrected *p*-values (*p*_FDR_) are reported for each region.

| **Region** | **Proportion mediated** | **95% confidence interval** | ***p*_FDR_** |
| --- | --- | --- | --- |
| Cortical gray matter | -0.322 | -1.032 – -0.063 | 0.024 |
| White matter | -0.417 | -1.506 – -0.099 | 0.024 |
| Deep gray matter | -0.288 | -0.939 – -0.075 | 0.024 |
| Total brain volume | -0.391 | -1.436 – -0.107 | 0.024 |
| Intracranial volume | -0.351 | -1.058 – -0.089 | 0.024 |
| Left hippocampus | -0.219 | -0.739 – -0.042 | 0.026 |
| Left amygdala | -0.261 | -0.946 – -0.052 | 0.006 |
| Right amygdala | -0.254 | -0.875 – -0.042 | 0.014 |
| Left parahippocampal gyrus (anterior part) | -0.211 | -0.815 – -0.034 | 0.034 |
| Right parahippocampal gyrus (anterior part) | -0.160 | -0.508 – -0.033 | 0.024 |
| Medial and inferior temporal gyri anterior part left | -0.211 | -0.731 – -0.035 | 0.024 |
| Medial and inferior temporal gyri anterior part right | -0.222 | -0.743 – -0.059 | 0.024 |
| Lateral occipitotemporal gyrus – fusiformis anterior part left | -0.146 | -0.466 – -0.018 | 0.034 |
| Lateral occipitotemporal gyrus – fusiformis anterior part right | -0.150 | -0.478 – -0.011 | 0.041 |
| Cerebellum right | -0.201 | -0.767 – -0.010 | 0.041 |
| Insula right | -0.352 | -1.124 – -0.099 | 0.024 |
| Insula left | -0.374 | -1.249 – -0.118 | 0.024 |
| Occipital lobe right | -0.204 | -0.643 – -0.022 | 0.039 |
| Medial and inferior temporal gyri posterior part right | -0.363 | -1.234 – -0.068 | 0.026 |
| Medial and inferior temporal gyri posterior part left | -0.295 | -0.994 – -0.064 | 0.024 |
| Cingulate gyrus – posterior part right | -0.250 | -0.846 – -0.052 | 0.024 |
| Cingulate gyrus – posterior part left | -0.197 | -0.675 – -0.036 | 0.024 |
| Frontal lobe left | -0.290 | -0.980 – -0.043 | 0.026 |
| Parietal lobe left | -0.288 | -0.974 – -0.055 | 0.024 |
| Caudate nucleus right | -0.215 | -0.716 – -0.049 | 0.024 |
| Caudate nucleus left | -0.166 | -0.671 – -0.027 | 0.024 |
| Thalamus right | -0.196 | -0.659 – -0.042 | 0.024 |
| Thalamus left | -0.238 | -0.867 – -0.055 | 0.024 |
| Lentiform Nucleus left | -0.243 | -0.908 – -0.010 | 0.041 |

**Supplementary Table 14. Brain volumes mediating the association between gestational age at birth and language outcomes**

Values represent the proportion of the total effect of gestational age at birth on language outcomes mediated by brain volumes along with associated confidence intervals (95%) and FDR-corrected *p*-values.

| **Region** | **Proportion mediated** | **95% confidence interval** | ***p*_FDR_** |
| --- | --- | --- | --- |
| Total brain volume | 0.320 | -0.120 – 0.849 | 0.225 |
| Total cortical grey matter | 0.377 | -0.045 – 0.869 | 0.199 |
| Total subcortical grey matter | 0.344 | -0.120 – 0.849 | 0.199 |
| Total white matter | 0.155 | -0.043 – 0.402 | 0.199 |
| Hippocampus left | 0.187 | -0.043 – 0.489 | 0.199 |
| Amygdala left | 0.171 | -0.062 – 0.4660 | 0.238 |
| Amygdala right | 0.181 | -0.085 – 0.508 | 0.199 |
| Gyri parahippocampalis et ambiens anterior part left | 0.139 | -0.027 – 0.360 | 0.199 |
| Gyri parahippocampalis et ambiens anterior part right | 0.213 | -0.012 – 0.507 | 0.199 |
| Medial and inferior temporal gyri anterior part left | 0.356 | -0.045 – 0.866 | 0.199 |
| Medial and inferior temporal gyri anterior part right | 0.421 | -0.055 – 0.943 | 0.199 |
| Lateral occipitotemporal gyrus – fusiformis anterior part left | 0.296 | -0.016 – 0.729 | 0.199 |
| Lateral occipitotemporal gyrus – fusiformis anterior part right | 0.295 | -0.041 – 0.763 | 0.199 |
| Cerebellum right | 0.413 | -0.055 – 1.240 | 0.238 |
| Insula right | 0.325 | -0.053 – 0.801 | 0.199 |
| Insula left | 0.293 | -0.042 – 0.760 | 0.199 |
| Occipital lobe right | 0.340 | -0.126 – 0.900 | 0.199 |
| Medial and inferior temporal gyri posterior part right | 0.322 | -0.185 – 0.892 | 0.238 |
| Medial and inferior temporal gyri posterior part left | 0.281 | -0.142 – 0.846 | 0.238 |
| Cingulate gyrus – posterior part right | 0.210 | -0.142 – 0.543 | 0.199 |
| Cingulate gyrus – posterior part left | 0.265 | -0.058 – 0.725 | 0.199 |
| Frontal lobe left | 0.311 | -0.194 – 0.968 | 0.240 |
| Parietal lobe left | 0.499 | -0.126 – 1.198 | 0.199 |
| Caudate nucleus right | 0.275 | 0.040 – 0.629 | 0.199 |
| Caudate nucleus left | 0.243 | 0.040 – 0.629 | 0.199 |
| Thalamus right | 0.282 | -0.118 – 0.767 | 0.237 |
| Thalamus left | 0.341 | -0.050 – 0.882 | 0.199 |
| Lentiform Nucleus left | 0.224 | -0.165 – 0.679 | 0.271 |

**Supplementary Table 15. Brain volumes mediating the association between gestational age at birth and motor outcomes after controlling for total brain volume**

Values represent the proportion of the total effect of gestational age at birth on motor outcomes mediated by brain volumes along with associated confidence intervals (95%) and FDR-corrected *p*-values.

| **Region** | **Proportion mediated** | **95% confidence interval** | ***p*_FDR_** |
| --- | --- | --- | --- |
| Brainstem | -0.106 | -5.381 – 4.330 | 0.924 |
| Left cerebellum | -0.567 | -16.152 – 2.030 | 0.924 |

**Supplementary Table 16. Regional brain volumes mediating the association between birth weight and cognitive outcomes**

Values represent the proportion of the total effect of birth weight on cognitive outcomes mediated by brain volumes along with associated confidence intervals (95%) and FDR-corrected *p*-values.

| **Region** | **Proportion mediated** | **95% confidence interval** | ***p*_FDR_** |
| --- | --- | --- | --- |
| Cortical grey matter | 0.758 | 0.290 – 2.33 | 0.04 |
| Right occipital lobe | 0.539 | 0.088– 1.18 | 0.04 |
| Medial and inferior gyrus | 0.494 | 0.021 – 1.12 | 0.04 |
| Left thalamus | 0.419 | 0.034 – 1.02 | 0.04 |
| Right parahippocampal gyrus (anterior part) | 0.233 | 0.035 – 0.50 | 0.04 |
| Left lateral occipitotemporal gyrus | 0.318 | 0.021 – 0.750 | 0.04 |

**Table 17. Regional brain volumes mediating the association between birth weight and language outcomes**

Values represent the proportion of the total effect of birth weight on language outcomes mediated by brain volumes along with associated confidence intervals (95%) and FDR-corrected *p*-values.

| **Region** | **Proportion mediated** | **95% confidence interval** | ***p*_FDR_** |
| --- | --- | --- | --- |
| Cortical gray matter | 0.758 | 0.290 – 3.153 | 0.018 |
| White matter | 0.479 | 0.056 – 1.729 | 0.026 |
| Cerebellum | 0.665 | 0.249 – 2.444 | 0.018 |
| Deep Gray Matter | 0.737 | 0.255 – 2.654 | 0.018 |
| Total | 0.747 | 0.296 – 2.785 | 0.018 |
| ICV | 0.826 | 0.250 – 3.394 | 0.018 |
| Left hippocampus | 0.341 | 0.067 – 1.739 | 0.019 |
| Left amygdala | 0.435 | 0.095 – 2.005 | 0.018 |
| Right amygdala | 0.464 | 0.094 – 1.722 | 0.021 |
| Left parahippocampal gyrus (anterior part) | 0.266 | 0.058 – 1.032 | 0.018 |
| Right parahippocampal gyrus (anterior part) | 0.343 | 0.104 – 1.286 | 0.018 |
| Medial and inferior temporal gyri anterior part left | 0.600 | 0.215 – 2.034 | 0.018 |
| Medial and inferior temporal gyri anterior part right | 0.721 | 0.289 – 2.626 | 0.018 |
| Lateral occipitotemporal gyrus – fusiformis anterior part left | 0.480 | 0.164 – 1.938 | 0.018 |
| Lateral occipitotemporal gyrus – fusiformis anterior part right | 0.511 | 0.167 – 1.775 | 0.018 |
| Cerebellum right | 0.679 | 0.304 – 2.406 | 0.018 |
| Insula right | 0.605 | 0.237 – 2.301 | 0.000 |
| Insula left | 0.582 | 0.221 – 2.014 | 0.018 |
| Occipital lobe right | 0.670 | 0.225 – 2.076 | 0.018 |
| Medial and inferior temporal gyri posterior part right | 0.751 | 0.237 – 3.115 | 0.018 |
| Medial and inferior temporal gyri posterior part left | 0.649 | 0.263 – 2.374 | 0.018 |
| Cingulate gyrus – posterior part right | 0.575 | 0.172 – 2.265 | 0.018 |
| Cingulate gyrus – posterior part left | 0.671 | 0.238 – 2.492 | 0.018 |
| Frontal lobe left | 0.686 | 0.265 – 2.995 | 0.018 |
| Parietal lobe left | 0.841 | 0.308 – 3.229 | 0.018 |
| Caudate nucleus right | 0.488 | 0.154 – 1.767 | 0.018 |
| Caudate nucleus left | 0.466 | 0.155 – 1.796 | 0.018 |
| Thalamus right | 0.626 | 0.191 – 2.153 | 0.018 |
| Thalamus left | 0.698 | 0.217 – 2.528 | 0.019 |
| Lentiform Nucleus left | 0.561 | 0.185 – 1.882 | 0.018 |
